# Development, dimorphism, and divergence of the oral dentition in threespine sticklebacks (*Gasterosteus aculeatus*)

**DOI:** 10.64898/2026.05.15.725320

**Authors:** Adriana G. Mendizabal, Craig T. Miller

## Abstract

How morphology forms during development and changes during evolution remain major questions in biology. In vertebrates, teeth have long served as model systems to address these questions. In threespine stickleback fish (*Gasterosteus aculeatus*), repeated and convergent increases in pharyngeal tooth number in derived freshwater sticklebacks occur, suggesting increased tooth number is adaptive in freshwater environments, likely due to a diet of larger prey in freshwater. Whether changes in oral tooth patterning also occur in freshwater sticklebacks was unknown. Here we describe oral tooth number and patterning in a dense developmental time course of lab-reared ancestral marine and derived freshwater fish. We address three major questions. First, is the spatial sequence of early oral tooth formation invariant as we previously described for the pharyngeal dentition? Second, is oral tooth patterning in the upper and lower jaw sexually dimorphic, and if so, when during development does this dimorphism arise? Third, have freshwater fish evolved increases in oral tooth number? We find that (1) unlike the pharyngeal dentition, the oral jaw early spatial sequence is variable, especially in the lower jaw (2) sexual dimorphism in both oral jaws arises at the late juvenile stage with males having more teeth and (3) freshwater fish have evolved more oral teeth similar to the evolved tooth gain in the pharyngeal jaw. Together our morphological descriptions advance the stickleback oral jaw as a model system to study how morphology forms during development and evolves in nature.

## INTRODUCTION

Most non-mammalian vertebrates, including fish, have retained the ancestral condition of polyphyodonty, continuous tooth replacement throughout the lifespan^1–5^. Still, the mechanisms involved in the formation and replacement of teeth and other epithelial appendages are largely homologous between non-mammalian and mammalian vertebrates^3,5,6,7^. As such, tooth formation and regeneration are attractive models to study vertebrate organogenesis and morphogenesis. The diversity of tooth patterning in bony fish (teleosts) makes fish a powerful model for answering questions about the patterning and sequence of early tooth development and replacement^8–14^.

The presence of oral teeth is not universal among teleosts. For example, Cyprinids such as zebrafish (*Danio rerio*) possess only pharyngeal teeth, while other teleosts such as threespine sticklebacks (*Gasterosteus aculeatus*) have both pharyngeal and oral teeth^15–17^. Oral teeth are thought to be used for prey capture and nest building, while pharyngeal teeth seem to primarily be used for prey mastication^18–21^. Despite the difference in biological function of the two dentitions, oral and pharyngeal teeth form through similar gene expression patterns and share many morphological characteristics^15,22–24^. In sticklebacks, despite similar genes being involved in oral and pharyngeal tooth replacement, the mechanisms by which they occur might differ. While in pharyngeal teeth, replacement seems to typically involve the dislodging of the old tooth; in the oral jaw, the replacement tooth typically appears before shedding of the replaced tooth^25^.

To better understand early tooth development and replacement, we previously described the primary tooth sequence of the pharyngeal dentition, finding that the first 17 pharyngeal teeth form in a stereotypical spatial sequence before the first replacement event of the pioneer tooth^15^. However, little is known about the oral dentition, both with respect to how it forms as well as when the first replacement event is initiated.

The oral dentition in the upper jaw (premaxilla) was previously shown to be variably sexually dimorphic in wild marine and freshwater populations, with males having more teeth^26^. However, three questions regarding this sexual dimorphism in oral tooth patterning remain: (1) whether this sexual dimorphism is genetic and present in lab-reared populations fed the same food (2) if so, when during development this sexual dimorphism arises, and (3) whether sexual dimorphism in tooth patterning is also present in the lower jaw.

Given that the selective pressures following adaptive radiation affect both sexes uniformly, it has been hypothesized that sex-specific selection might weaken in populations that are adapting to new environments^27–30^. Recent work has provided support for this divergence hypothesis, showing that sexual dimorphism in traits such as body shape, pelvic girdle length, and gill raker length were reduced in freshwater populations when compared to their marine ancestors^31–35^. Whether sexual dimorphism is also reduced upon freshwater adaptation for oral tooth patterning was unknown.

Further, the evolutionary history of sticklebacks, having arisen in a marine environment and repeatedly colonized freshwater environments, has led to an array of divergent traits between ancestral marine and derived freshwater adults, including a reduction in lateral plate armor, loss of pelvic fins, reduced dorsal spine length, and an increase in pharyngeal tooth number in freshwater populations^36–40^. However, less is known about how the oral dentition differs between the ancestral marine and derived freshwater populations. Here we present a detailed morphological characterization of oral tooth development in both marine and freshwater populations to test whether differences in oral tooth patterning have evolved as well. By comparing oral tooth number between populations throughout juvenile and adult stages we also test whether the spatial sequence of primary oral tooth formation is invariant and whether sexual dimorphism is present in lab-reared stocks. Together, these data provide a more comprehensive understanding of how the morphology of the oral dentition forms during development and evolves in nature.

## RESULTS

We previously described the development of the pharyngeal dentition in lab-reared sticklebacks and found that the early sequence of tooth formation was largely invariant and followed a consensus sequence until the first tooth replacement event^15^. Here, we first tested whether the spatial sequence of early oral tooth development is also invariant through the collection of dense developmental time courses of both marine and freshwater lab-reared fish.

In sticklebacks, oral teeth are present on both the upper jaw (premaxilla) and lower jaw (dentary)^41^. Both the jaw and teeth can be visualized by staining with Alizarin red, which labels bone and teeth fluorescently red (**Figure 1A-B**). In our developmental time course, we found that in both marine and freshwater populations, oral teeth do not begin mineralizing until soon after hatching, with a pioneer tooth emerging on both the premaxilla and dentary between 10-11 days post fertilization (dpf), 2-3 days after hatching (**Figure 2A,E**). In both marine and freshwater populations, a second tooth emerged on the premaxilla and dentary between 12-13 dpf (**Figure 2B,F**). While the onset of second tooth formation was consistent across populations, we observed surprising differences in the degree of variability in second tooth positioning. Namely, the positioning of the second tooth was more variable in marine than freshwater populations. Specifically, in marine fish, the second tooth emerged medial to the pioneer in 64% of instances (n=7/11) and lateral in the remaining 36% (n=4) (**Figure S1A**). By contrast, in freshwater populations, the position of the second tooth was highly stereotypical, emerging medial to the pioneer in 94% of instances (n=15/16) (**Figure S1A**). Notably, unlike in the premaxilla, second-tooth positioning on the dentary was divergent across populations. In 79% of marine fish, the second tooth was positioned medially to the pioneer tooth (n=11/14, **Figure S1B**), while in freshwater fish, it was most often positioned lateral to the pioneer (n=10/15, **Figure S1B**). Thus, the spatial sequence of early tooth formation in the oral dentition in lab-reared sticklebacks is strikingly more variable than in the pharyngeal dentition, with variation in the spatial patterning of the oral jaw present by the two-tooth stage, unlike the early largely invariant sequence of the first 17 pharyngeal teeth we previously described^15^.

**Figure 1.**
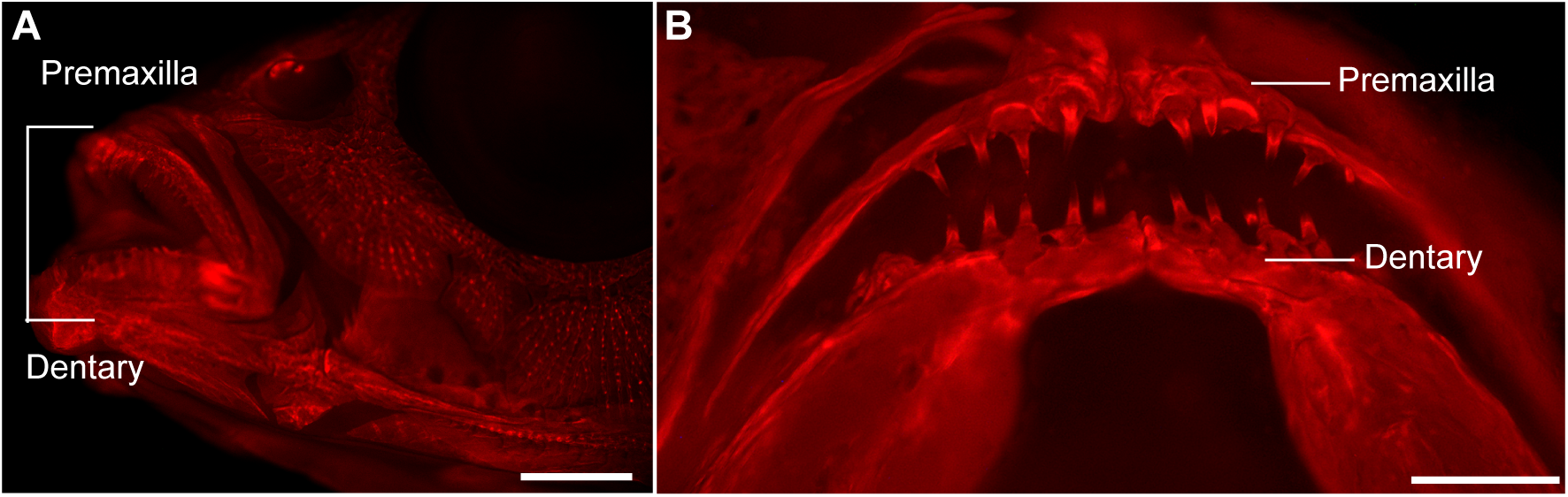
Threespine stickleback (*Gasterosteus aculeatus*) oral jaw. (A) Adult freshwater stickleback stained with Alizarin red in whole mount to label mineralized bone and imaged under fluorescence. (B) Anterior view of upper jaw (premaxilla) and lower jaw (dentary) from a 41 days post fertilization (dpf) freshwater fish larva. Scale bars = 250 µm (A), 100µm (B).

**Figure 2.**
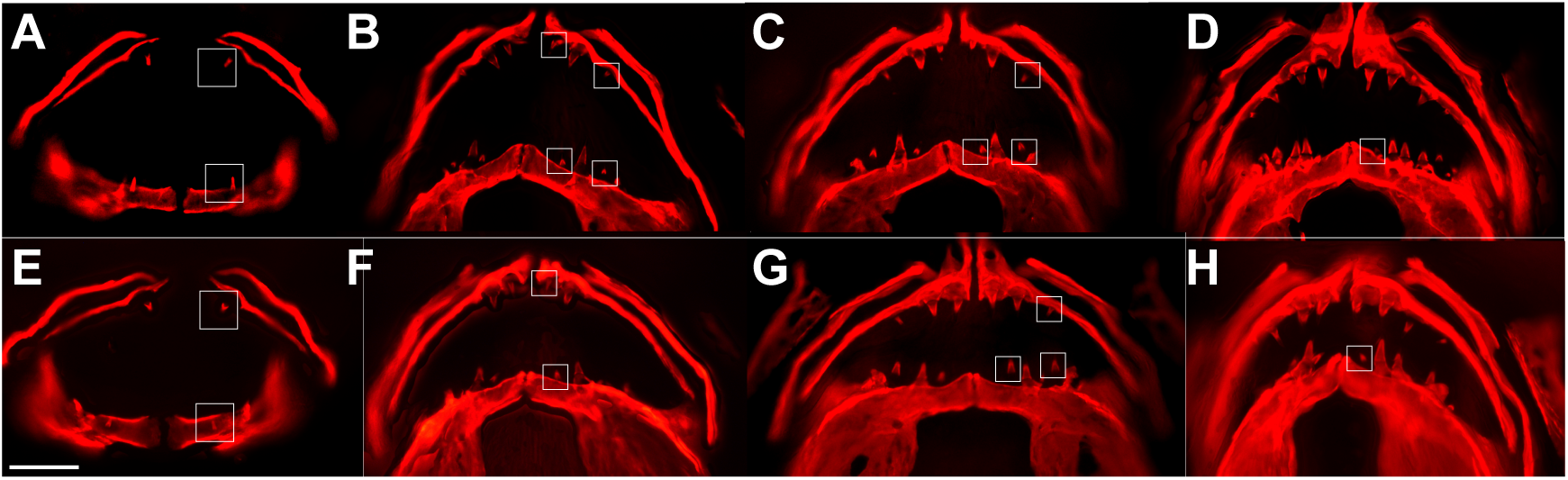
Developmental time course of oral tooth formation in marine and freshwater threespine sticklebacks. Anterior view of representative time course images from marine (A-D) and freshwater (E-H) fish larvae at 10-11 dpf (A,E); 12-13 dpf (B,F), 13-14 dpf (C,G); and 15-17 dpf (D,H) stages. Jaws are dissected and stained with Alizarin red to label mineralized bone. White boxes indicate newly forming mineralized teeth. Scale bar = 100 µm; dpf = days post fertilization.

To test whether differences in tooth-positioning variability persisted between marine and freshwater populations, we assessed the positioning and timing of third-tooth initiation. In the marine population, we found that the third tooth most often emerged between 13-14 dpf on both the premaxilla and dentary (n=9; **Figure 2C**; **Figure S2-5**). Relative to their marine counterparts, the timing of third tooth formation in the premaxilla of freshwater fish displayed greater variability, with roughly equal numbers of fish forming three teeth at the 13-14 dpf (n=12) and 15-17 dpf (n=14) stages (**Figure 2G; Figure S2,S4**). Similarly, we observed greater temporal variability in the dentary, with 28% of fish forming a third tooth by 11-12 dpf (n=8), 45% of fish forming a third tooth between 13-14 dpf (n=13), and the remainder of fish forming a third tooth between 15-17 dpf (n=8) (**Figure S3,S5**). Despite the variability in timing of tooth emergence, when three teeth were present in either jaw, the pioneer was in the center. Similarly, in both populations, the distance between the pioneer and tooth most medial to it was consistently shorter than the distance between the pioneer and tooth in the lateral position (**Figure S6**).

We collected later time points in both populations to assess the degree to which spatiotemporal patterning was conserved beyond the third-tooth stage. In marine populations, we found that a fourth premaxillary tooth first emerged between 13-14 dpf (n=3); however, the majority of fish displayed a fourth tooth at 15-17 dpf (n=5, **Figure 2D**). In contrast in this time course, freshwater fish did not display evidence of a developing fourth tooth until the 16-17 dpf stage (**Figure 2H)**.

Interestingly, despite this temporal variation, the emergence of the fourth premaxillary tooth was spatially consistent across both populations, forming in the position most lateral to the pioneer and on the same spatial plane as the first three teeth (**Figure 2D,H**; n=10). In the dentary of marine fish, the fourth tooth was similarly positioned lateral to the third tooth (position 4) and appeared on the same spatial plane as the first three teeth. Thus, in marine populations, the positioning of the first four primary teeth is conserved between jaws. Interestingly, we found that in freshwater populations, the position of a fourth tooth on the dentary was more variable, emerging in one of three positions: lateral to the second tooth (position 4, n=4); medial to the third tooth (position 5, n=4); and in between the pioneer and third tooth (position 6, n=4) (**Figure S5**).

Notably, in both populations, when the fourth tooth emerged on the dentary at position 4, it appeared to be on the same plane as the first three teeth; while when it emerged in positions 5 and 6, it appeared to be on a second spatial plane (**Figure 3A,D)**. Given that the initiation of a second row is not contingent on the number of preexisting teeth, this spatial shift is likely not due to a spatial constraint on the jaw.

**Figure 3.**
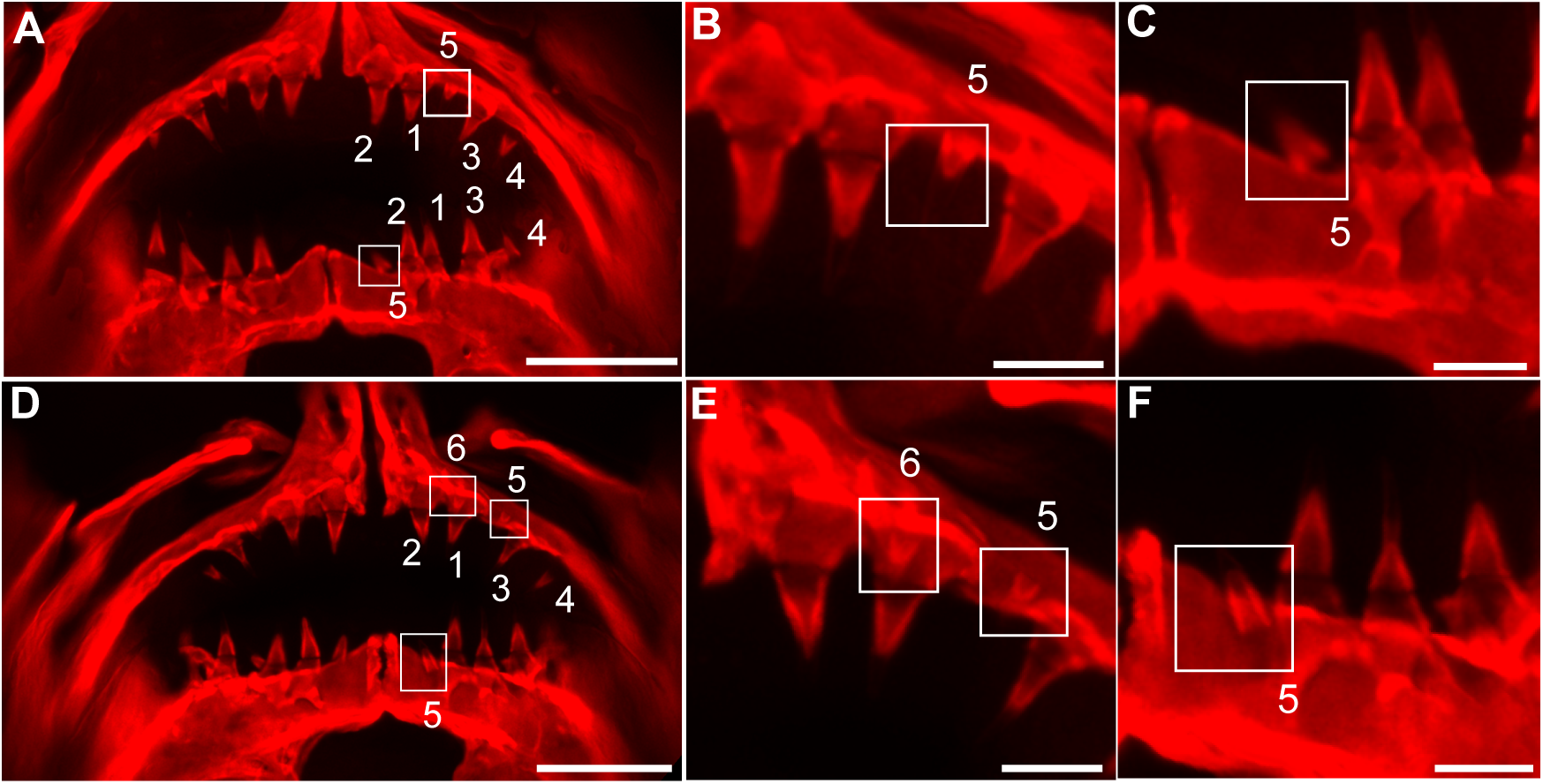
Early spatial patterning in the primary oral dentition. Anterior view of representative Alizarin-stained marine (A-C) and freshwater (D-F) stickleback fish from 18-20 days post fertilization (dpf). Numbers 1-4 denote the first four primary teeth forming in the same spatial plane. The white boxes indicate the beginning of a second tooth row, initiated in positions 5 and 6. Higher magnification views show tooth initiation on the premaxilla (B,E) and dentary (C,F). N=12. Scale bars = 100 µm (A, D); 25 µm (B-C, E-F).

We continued following the sequence of oral tooth formation in both populations to determine whether all emerging mineralized baby teeth after the four-tooth stage formed on the second row. Ultimately, we found that in both populations, a fifth tooth emerged on the premaxilla between 18-20 dpf; was positioned between the pioneer and third tooth (position 5); and began the formation of a second tooth row (n=12, **Figure 3B,E**). Similarly, in both populations, a fifth tooth emerged only after the first three teeth became ankylosed (**Figure S2**, **Figure S2-S3**). Across both populations, the timing of when a fifth tooth emerged on the dentary was wider, ranging from 13-20 dpf to 18-20 dpf (n=18, **Figure S4**, **Figure S4-S5**). However, in both populations, the fifth tooth was most often positioned medial to the pioneer (position 5) and formed on a second tooth row (**Figure 3C,F**). In both populations, the emergence of a sixth tooth occurred within the same developmental time window as the fifth tooth and was positioned between the first and third primary teeth (**Figure 3D-E**). Given the previously described divergence in second tooth positioning between marine and freshwater populations, the conserved positioning of the sixth tooth (on a second spatial plane between the pioneer and tooth most medial to it, irrespective of whether that medial tooth appeared second or third in the sequence), suggests that this position is more developmentally constrained.

Notably, positions 5 and 6 were occupied at earlier developmental stages in freshwater populations than in marine populations (**Figures S2-S5**). Further, given our earlier observations about the spatial arrangement of teeth, we sought to determine whether subsequent teeth continued to form on the second row. To this end, we collected later time points and found that neither premaxillary nor dentary tooth number increased significantly between the 18-20 dpf and 25-28 dpf stages (**Figure 4A-F**). Specifically, the majority of both marine and freshwater fish displayed between five and six teeth on the premaxilla and dentary (**Figure S7**). However, we noted slightly greater variability in freshwater fish, with several individuals having 7-8 teeth on the premaxilla and dentary (**Figure S7 B,D**). Despite the slowed rate of new tooth formation between the 18-20 dpf and 25-28 dpf stages, the maturation of existing teeth increased substantially, with 60-80% of fish from 25-28 dpf having between 5-7 teeth ankylosed (n=17, **Figures S2-S5**).

**Figure 4.**
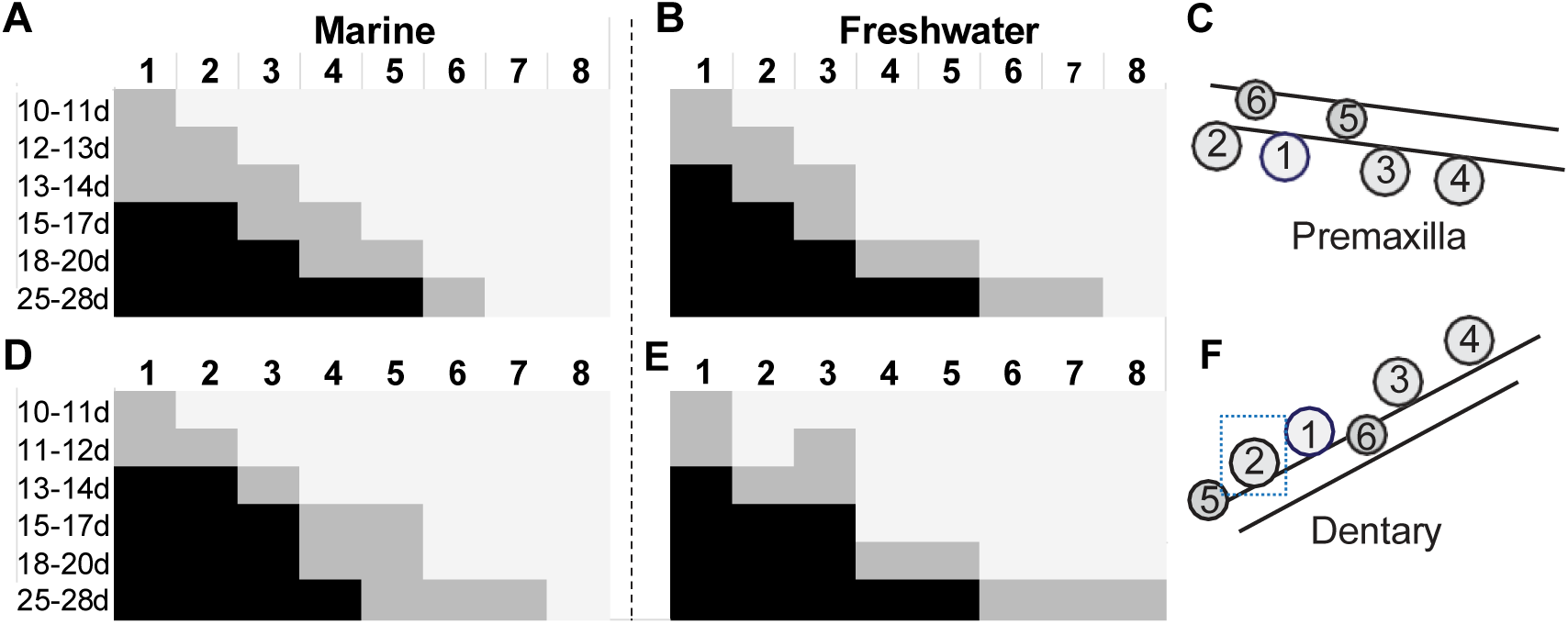
Consensus sequence of primary tooth positions on the premaxilla and dentary of marine and freshwater populations. (A-B, D-E) Numbered vertical rows indicate the position on the jaw. Horizontal numbers show the developmental time (dpf) that the observed sequence is present in the majority of fish. (A-C) Premaxillary consensus sequence for (A) marine and (B) freshwater populations. (C) Schematic depicting tooth positions on the premaxilla when viewed from the anterior position. (D-E) Dentary consensus sequence for (D) marine and (E) freshwater populations. (F) Schematic depicting tooth positions on the dentary for the ancestral marine population viewed from the anterior position. The dashed box highlights the second tooth variability in freshwater dentary positioning. Each schematic represents the left side of the fish. Colors of boxes indicate the absence of a tooth (light grey), presence of a developing germ (dark grey), or presence of an ankylosed tooth (black).

### Tooth maturation as a signal for the formation of new teeth

Interestingly, the relationship between the stage of preexisting mineralized baby teeth and the formation of new teeth seemed to shift across development. Specifically, in both populations, the formation of the first four teeth was relatively independent of the calcification stage of the preceding teeth (n=9; **Figure S2-S5**). However, the formation of the fifth and sixth teeth appeared to depend on the stage of calcification of the earlier teeth, with the fifth and sixth teeth forming only after the first three teeth become ankylosed (n=22; **Figure S2-S5**). Further, while the first four teeth appeared to form extraosseously, the fifth and sixth teeth exhibited evidence of intraosseous formation (**Figure 5A-B**). Altogether, in both populations, the first twelve days after hatching (∼18-20 dpf) are when the first five to six teeth form, with a tooth forming roughly every two days (**Figure 4A-B**; **D-E**). Given the positional variability observed between marine and freshwater fish early in development by the six-tooth stage, we did not attempt to continue to describe the spatial sequence by numbering subsequent oral tooth positions.

**Figure 5.**
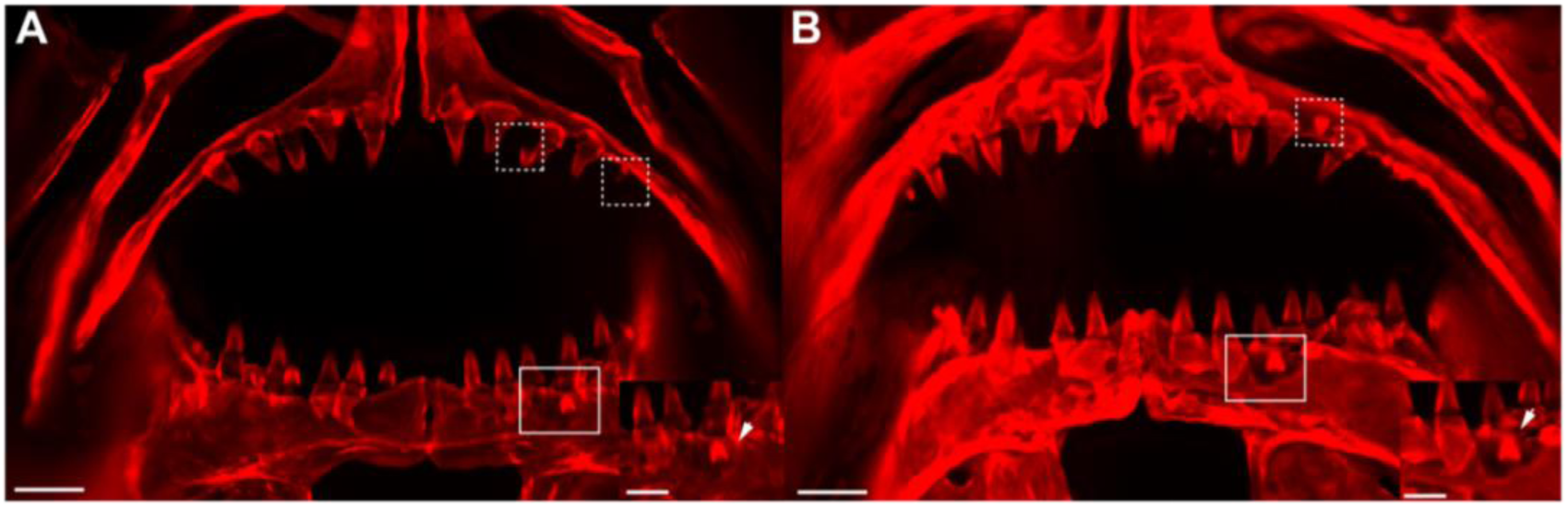
Intraosseous tooth formation in the oral jaw. Anterior views of Alizarin-red-stained, flat-mounted oral jaws of (A) marine and (B) freshwater fish. Representative images from the 25-28 dpf stage. Dashed boxes indicate newly forming teeth following apparent resorption. Solid boxes highlight teeth forming adjacent to the primary tooth. Scale bars = 50 µm; inset scale bars = 25 µm. dpf = days post fertilization.

### Upper-lower decoupling in the oral jaw

Given the positional and temporal variation observed in the primary tooth sequence of premaxillary and dentary tooth formation, we compared tooth number between the upper and lower jaws within individual fish to determine the prevalence of upper-lower jaw decoupling. We found that while left-right bilateral symmetry was maintained in the 92.1% of marine (n=70) and 97.4% of freshwater (n=74) fish, significant upper-lower jaw decoupling was present in both populations (**Figure 6A**). In marine fish, 43.4% of animals displayed decoupling in premaxillary and dentary tooth counts, while in freshwater populations, 48.7% of fish displayed decoupling between upper and lower jaws (**Figure 6A**). Interestingly, we observed a strong directional bias in upper-lower decoupling, with 84.8 and 89.2% of instances occurring when more teeth were present on the dentary in marine and freshwater populations, respectively (**Figure 6B**).

**Figure 6.**
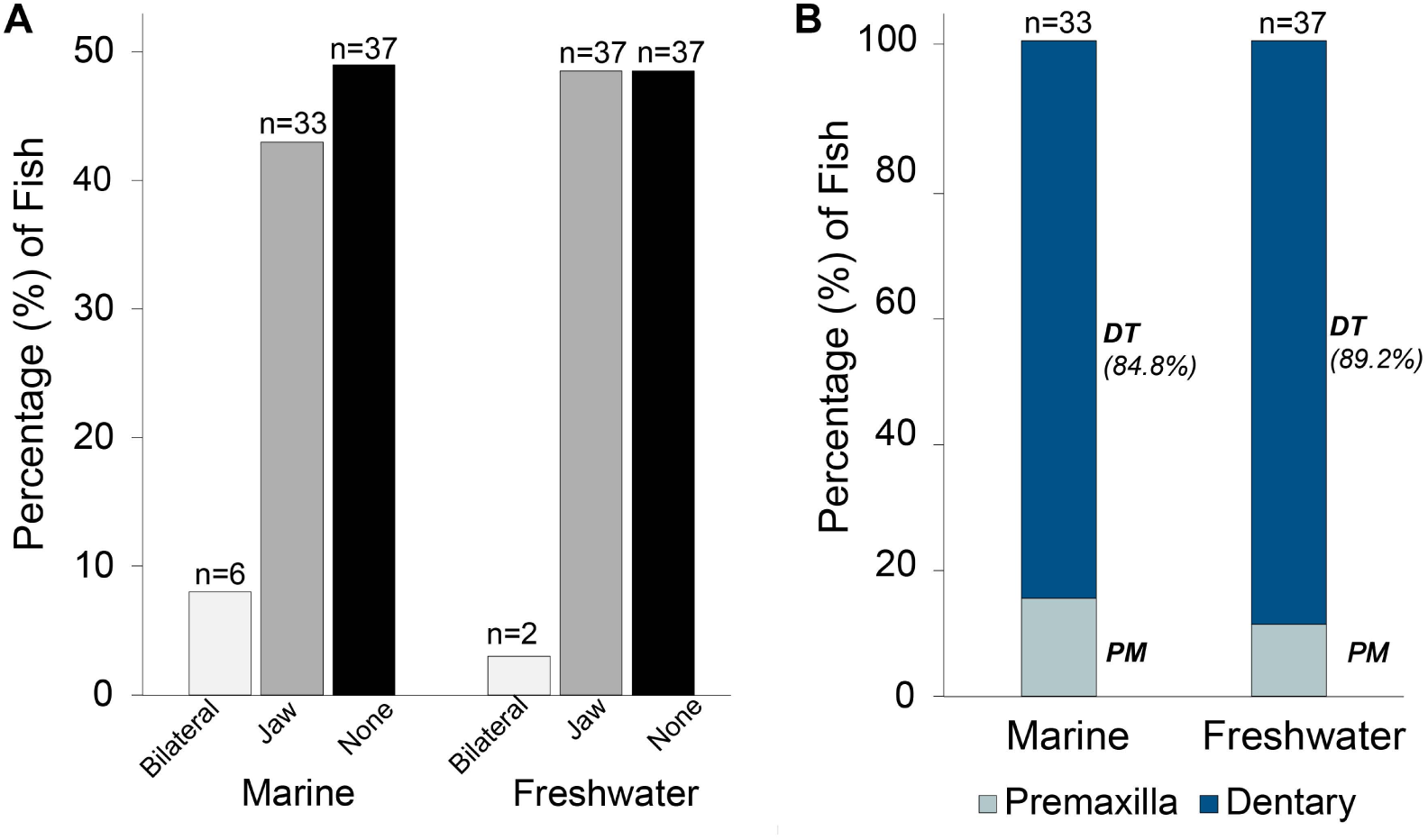
Oral jaw asymmetry and upper-lower jaw decoupling in marine and freshwater stickleback populations. (A) Bar graphs indicate percentage of animals exhibiting asymmetry/decoupling in tooth number between the left and right sides of the jaw (bilateral), premaxilla and dentary (jaw), and no asymmetry (none). Numbers above bars represent the number of individuals displaying that pattern. N = 152 (marine = 76; freshwater = 76). (B) Stacked bar graphs represent the directionality of asymmetry, with percentages indicating whether decoupling results in greater premaxillary or dentary tooth counts. N = 70 (marine n=33; freshwater n=37). Premaxilla (PM); Dentary (DT).

### Tooth shedding and replacement events on the oral jaw

We did not observe evidence of tooth shedding or replacement in either population by 25-28 dpf (**Figure S8**). We next collected later time points from 40-44 dpf freshwater fish to try to identify when the first replacement events occurred. At these stages, we observed several morphological indications of tooth shedding and potential replacement, including: (1) presence of resorption lacuna, large craters in the dental epithelium likely generated by osteoclast activity following tooth shedding^42–44^ (**Figure 7**, **Figure S9**), and (2) emerging newly mineralized teeth either within these craters or directly adjacent to primary teeth (**Figure 8**, **Figure S9**). Interestingly, these morphological indicators were observed on the dentary prior to the premaxilla (**Figure S9**), suggesting that the increased tooth formation rate in the dentary may relate to an earlier onset of replacement in the lower jaw than upper jaw. This earlier presence of signs of tooth replacement in the dentary than premaxilla is consistent with our earlier observations of the directional bias in upper-lower jaw decoupling, favoring higher tooth counts in the dentary.

**Figure 7.**
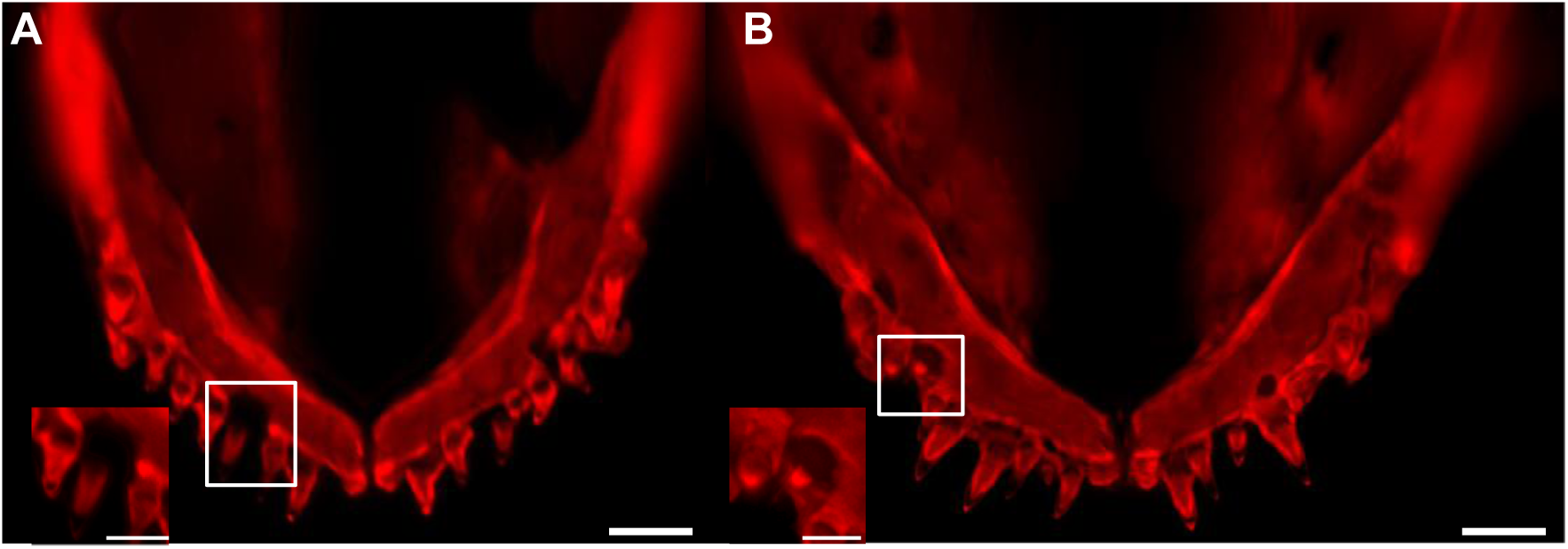
Morphological stages of lacunar resorption in the oral jaw. (A-B) Representative images of the Alizarin-red-stained dentary (dorsal view) of two freshwater sticklebacks from 40-43 dpf. (A) Evidence of a bony crater on the dentary that appears to have formed following osteoclast activity (n=6). (B) The emergence of a replacement tooth within the resorption crater. Scale bars = 100 µm, inset scale bars = 50 µm.

**Figure 8.**
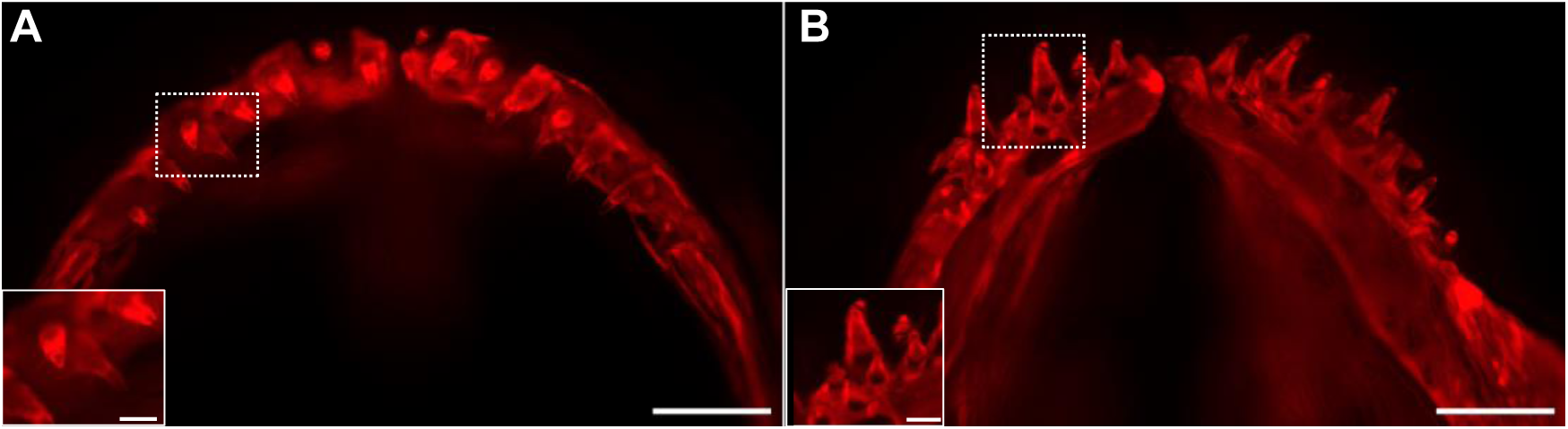
Formation of a multi-rowed dentition on the premaxilla and dentary of freshwater sticklebacks. (A-B) Representative Alizarin-red-stained images (dorsal view) highlighting the formation of a multi-rowed dentition on the premaxilla and dentary of a 41 dpf freshwater stickleback (n=10). Insets and boxes highlight the formation of a putative replacement tooth at the base of a primary tooth. Scale bars = 100 µm, inset scale bars = 25 µm.

### Sexual dimorphism in oral tooth number

Given the previously described report of sexual dimorphism in upper jaw premaxilla oral tooth number in wild stickleback populations^26^, we next sought to test whether sexual dimorphism is present in lab-reared fish. We quantified tooth number in marine and freshwater lab-reared populations from three developmental stages: young (<30 mm standard length), juvenile (30-40 mm), and adult (> 40 mm). Sticklebacks have a simple XY sex determination system^245^, and we genotyped the sex chromosomes in each fish as previously described^46^. We then assessed the degree of sexual dimorphism present across the entire developmental time range of ancestral marine and derived freshwater populations. In the marine population, we found a significant interaction between standard length (SL) and sex (*P* < 0.0001), driven by an increase in premaxillary tooth number in response to SL in males (*P* < 0.0001) and a non-significant relationship in females (*P* = 0.121; **Figure 9A**, **Table 1**). By contrast, in freshwater populations, SL significantly influenced premaxillary tooth number in both sexes (**Figure 9B**, **Table 2**).

**Figure 9.**
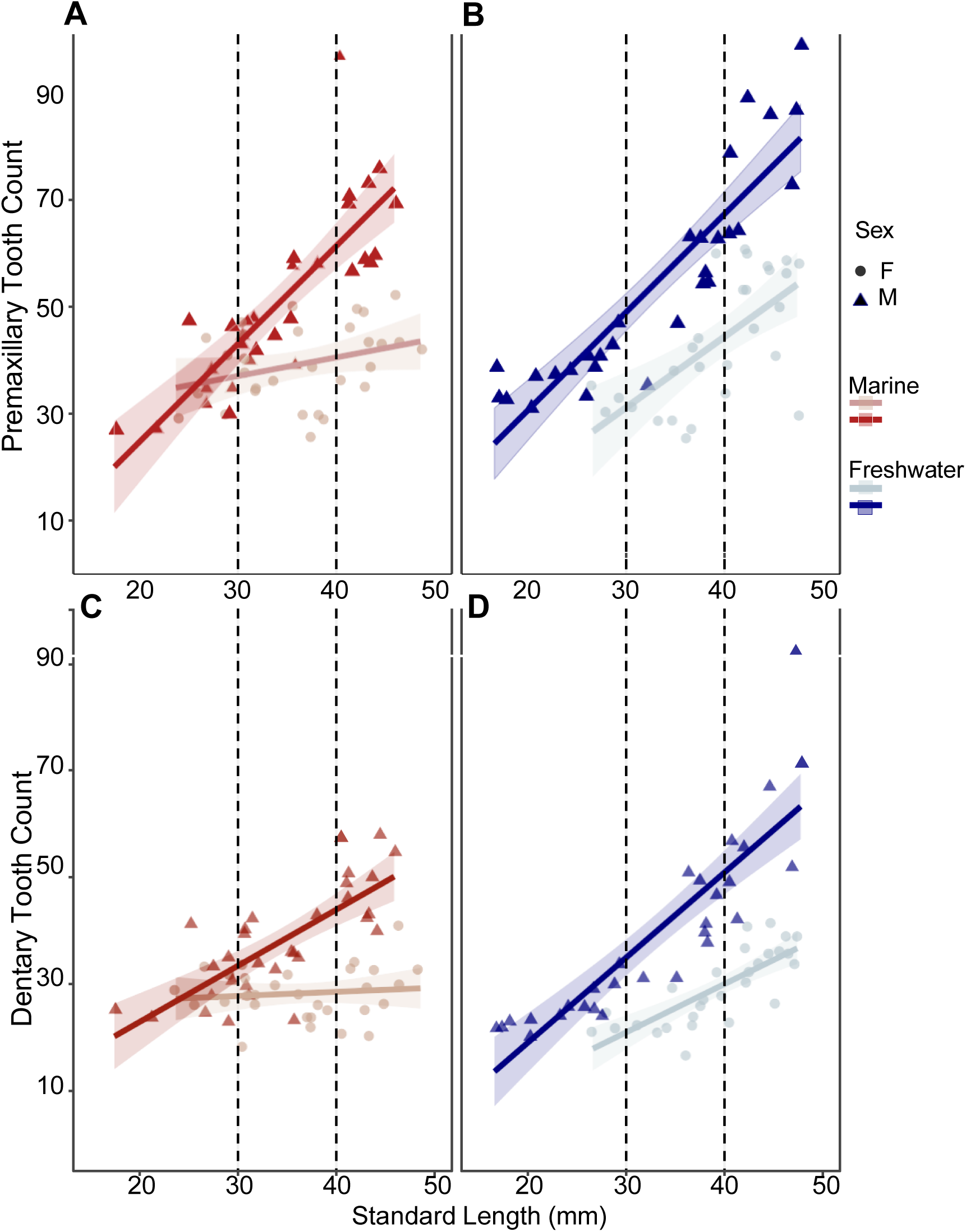
Sexual dimorphism in oral tooth number in marine and freshwater populations. (A-D) Linear regression of premaxillary (A-B) and dentary (C-D) tooth counts relative to standard length (SL) in marine (red) and freshwater (blue) populations. Points represent individual fish (triangles/dark lines = males; circles/light lines = females). Shaded regions indicate 95% confidence intervals; dashed lines denote developmental stage: young (< 30 mm), juvenile (30-40 mm), and adult (> 40 mm). Marine: n = 59, freshwater: n = 56.

**Table 1.** Summary of linear regression results for the relationship between tooth number and standard length in female and male marine sticklebacks. Regression estimates, standard errors, t-statistics, and *P*-values are shown for the relationship between tooth number in response to standard length in marine females and males. Slopes represent the rates of tooth addition per mm for each sex. Bolded *P*-values highlight the significant relationship between body size and tooth addition in male but not female sticklebacks. The significant interaction term (Male Slope Difference) indicates that tooth addition in response to increasing standard length is sex-dependent. Values in parentheses report 95% confidence intervals. The model fit is reported as R^2^ and adjusted R^2^ values. *P* values were determined using an Analysis of Covariance (ANCOVA). Statistical significance is denoted in bold. Total n=59 (F=30, M=29).

| *Predictors* | **Premaxilla Total** |  |  |  | **Dentary Total** |  |  |  |
| --- | --- | --- | --- | --- | --- | --- | --- | --- |
|  | *Estimates* | *std.*<br>*Error* | *Statistic* | *p* | *Estimates* | *std.*<br>*Error* | *Statistic* | *p* |
| Female Slope | 0.35<br>(-0.01 – 0.70) | 0.18 | 1.97 | 0.0542 | 0.08<br>(-0.22 – 0.37) | 0.15 | 0.52 | 0.6035 |
| Male Slope | 1.8<br>(-1.43 – 2.11) | 0.17 | 10.48 | **<0.0001** | 1.08<br>(-0.80 – 1.40) | 0.14 | 7.66 | **<0.0001** |
| Male Slope Difference (Interaction) | +1.42<br>(0.93 – 1.91) | 0.24 | 5.81 | **<0.0001** | +1.00<br>(0.59 – 1.41) | 0.20 | 4.91 | **<0.0001** |
| Observations | 59 |  |  |  | 59 |  |  |  |
| R<sup>2</sup> / R<sup>2</sup> adjusted | 0.731 / 0.716 |  |  |  | 0.642 / 0.622 |  |  |  |

**Table 2.** Summary of linear regression results for the relationship between tooth number and standard length in female and male freshwater sticklebacks. Regression estimates, standard errors, t-statistics, and *P* values are shown for the relationship between oral tooth number in response to standard length. Slopes represent the rates of tooth addition per mm for each sex. Bolded *P*-values highlight the significant relationship between body size and tooth addition in both sexes. The significant interaction term (Male Slope Difference) indicates that dentary tooth addition in response to increasing standard length is significantly different between the sexes. Values in parentheses report 95% confidence intervals. The model fit is reported as R^2^ and adjusted R^2^ values. *P* values were determined using an Analysis of Covariance (ANCOVA). Statistical significance is denoted in bold. Total n=56 (F=28, M=28).

| *Predictors* | **Premaxilla Total** |  |  |  | **Dentary Total** |  |  |  |
| --- | --- | --- | --- | --- | --- | --- | --- | --- |
|  | *Estimates* | *std. Error* | *Statistic* | *p* | *Estimates* | *std. Error* | *Statistic* | *p* |
| Female Slope | 1.32<br>(0.74 – 1.90) | 0.29 | 4.58 | **<0.0001** | 0.90<br>(0.56 – 1.25) | 0.17 | 5.25 | **<0.0001** |
| Male Slope | 1.80<br>(1.41 – 2.17) | 0.19 | 9.61 | **<0.0001** | 1.41<br>(1.18 – 1.63) | 0.11 | 12.64 | **<0.0001** |
| Male Slope Difference (Interaction) | +0.47<br>(-0.22 – 1.16) | 0.34 | 1.37 | 0.1767 | +0.50<br>(0.09 – 0.91) | 0.20 | 2.45 | **0.0176** |
| Observations | 56 |  |  |  | 56 |  |  |  |
| R<sup>2</sup> / R<sup>2</sup>adjusted | 0.712 / 0.695 |  |  |  | 0.808 / 0.797 |  |  |  |

Similarly, in the dentary of marine populations, the relationship between tooth number and standard length was influenced by sex, with a significant relationship between SL and tooth number in males, but not females (**Figure 9C**, **Table 1**). Interestingly, while in freshwater fish, dentary tooth number increased with SL in both sexes, males exhibited a significantly stronger relationship between tooth number and SL (**Figure 9D**, **Table 2**). Further, there was a higher degree of sexual dimorphism in the dentary than the premaxilla with respect to both overall tooth number and the rate of tooth increase relative to body size. Altogether, this finding supports what we observed earlier with respect to uncoupling between the development of the upper and lower jaws.

Given the significant interaction between sex and tooth number relative to body size described earlier, we assessed tooth number within each of the three developmental stages. Further, this analysis enabled closer examination into the onset of sexual dimorphism in tooth number. Our analysis revealed that sexual dimorphism in premaxillary tooth number was not observed in young fish from either marine or freshwater populations (**Figure 10A-B**). However, by the late juvenile stage, both marine and freshwater males possessed significantly more premaxillary teeth than females (Mar: +10.24; Fr: +18.32), with the difference being most pronounced in adults. Similarly, in the dentary, no significant differences in dentary tooth number were observed between young males and females from either population (**Figure 10C-D**). However, like the premaxilla, significant sexual dimorphism in the dentary was observed in the late juvenile stage (Mar: +8.06; Fr: +16.46) and was most pronounced in adults (Mar: +18.68; Fr: + 21.71, **Figure 10. C-D**). Given that length did not significantly impact tooth number within each developmental stage, raw tooth counts were used for statistical comparisons. Overall, the sexual dimorphism in oral tooth number appeared more severe in freshwater fish than marine fish (**Figures 9,10**), thus not supporting the hypothesis of reduced sexual dimorphism in derived freshwater populations.

**Figure 10.**
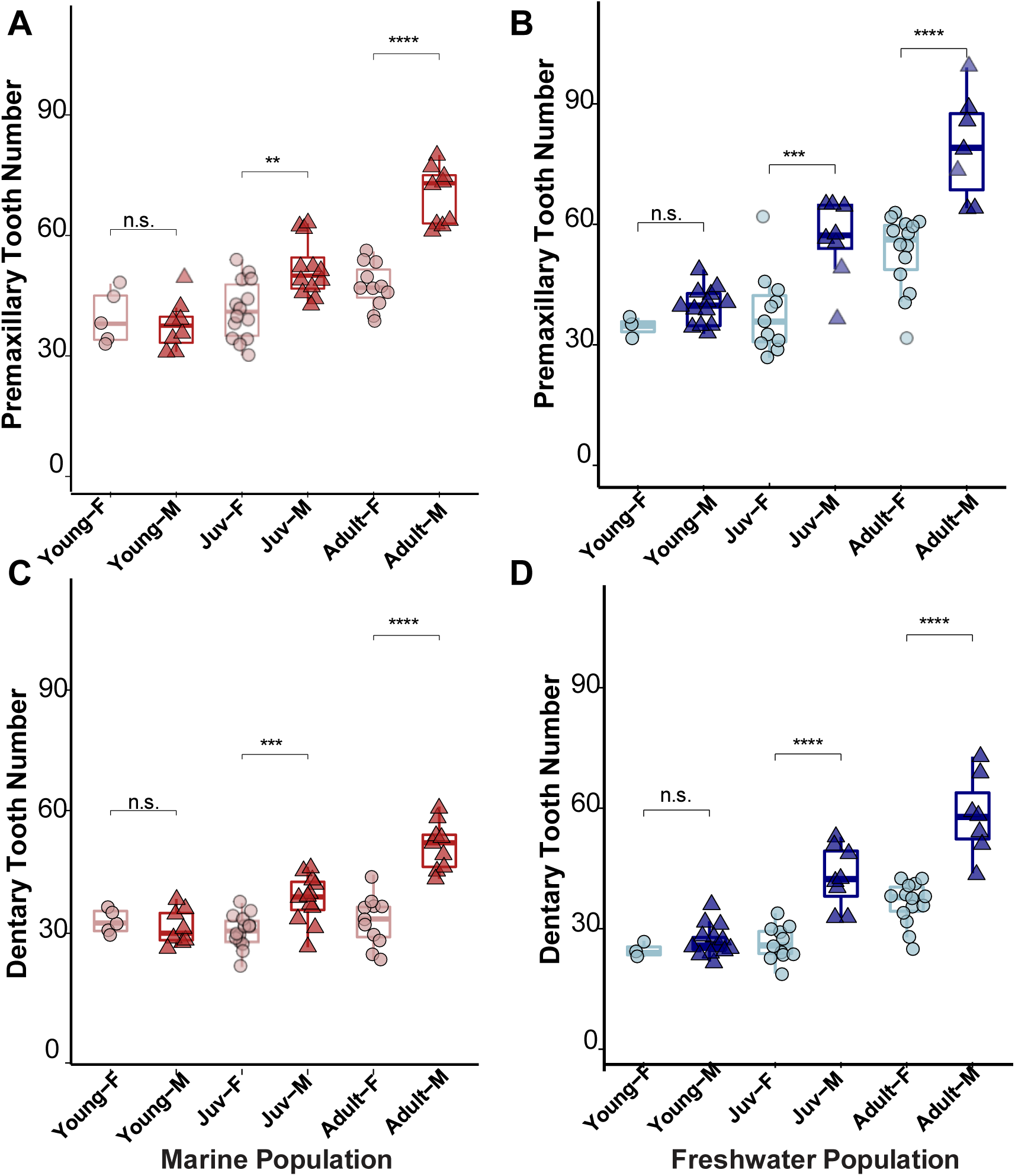
Sexual dimorphism in oral tooth number across discrete developmental stages: young (< 30 mm), juvenile (30-40 mm), and adult (> 40 mm). Premaxillary (A-B) and dentary (C-D) tooth counts in females and males from marine (A,C) and freshwater (B,D) populations. Boxes represent 25th-75th percentiles, midlines represent the median, whiskers denote boundaries of the min/max range. Standard length did not significantly impact tooth number within each developmental stage. Raw tooth counts were used for statistical comparisons. *P* values were determined using an Analysis of Covariance (ANCOVA). * *P*<0.05, \*\**P*<0.01, \*\*\**P*<0.001, \*\*\*\**P*<0.0001, n.s. not significant. Males = M, dark/triangles; Females = F, light/circles. Marine total: n = 59, freshwater total: n = 56. Marine: young n=13 (F=5, M=8); juvenile n=26 (F=14, M=12); adult n=20 (F=11, M=9). Freshwater: young n=16 (F=3, M=13); juvenile n=19 (F=11, M=8); adult n=21 (F=14, M=7).

### Evolved differences in oral tooth number between marine and freshwater populations

As previously described, a significant three-way interaction was observed between sex, standard length, and tooth number; wherein the relationship between standard length and tooth number was significant in both sexes in freshwater fish, but only in males in marine fish. This interaction was evident in how tooth number in females from each population responded to standard length. While tooth number in both jaws increased in response to standard length in freshwater, but not marine females, the difference was most pronounced in the dentary (**Figure 11**; **Table 3**).

**Figure 11.**
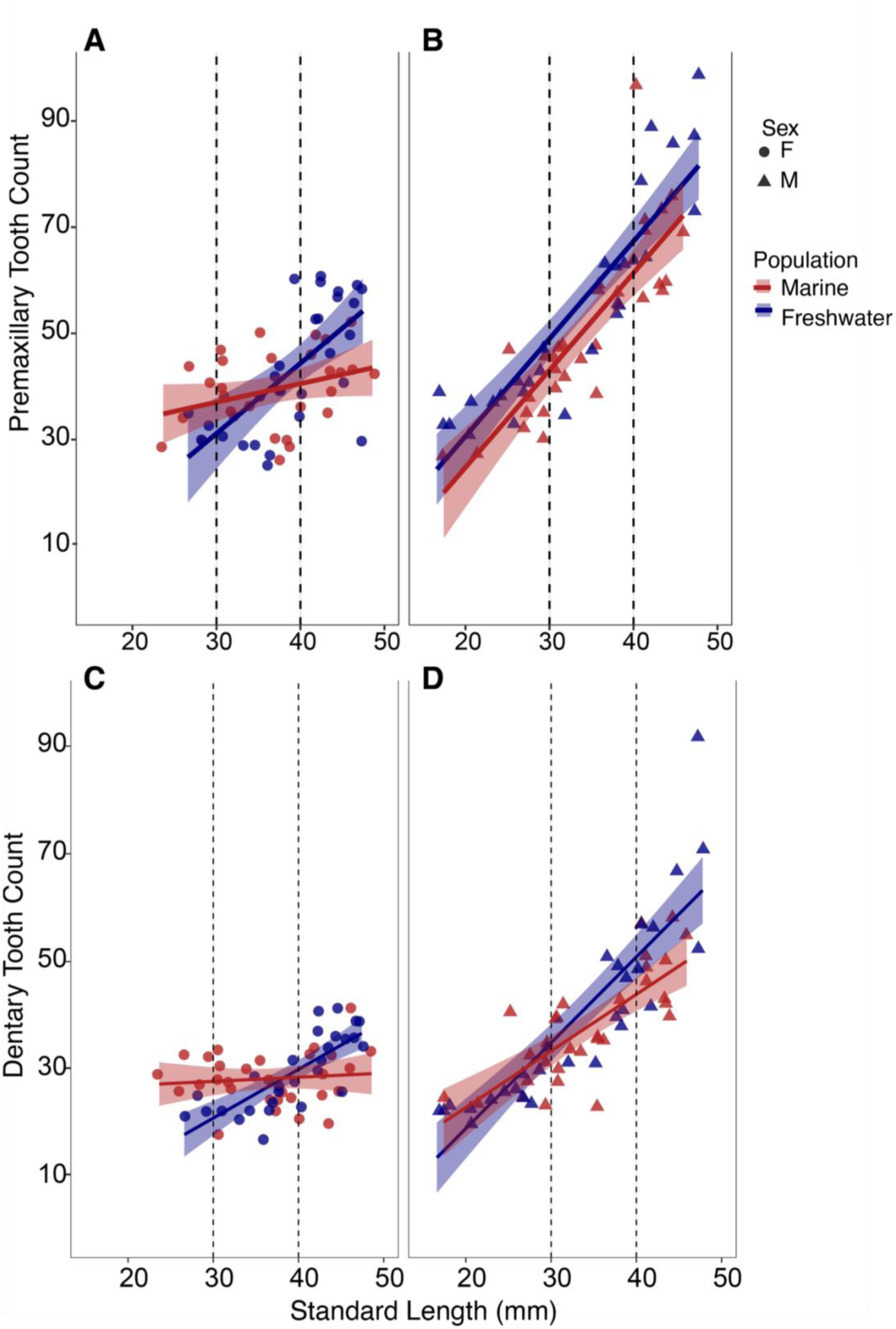
Population determines the effect of standard length on oral tooth number in females. Linear regression of (A-B) premaxillary and (C-D) dentary tooth counts relative to standard length in marine (red) and freshwater (blue) populations. Points represent individual fish (triangles = males; circles = females). Shaded regions indicate 95% confidence intervals; dashed lines denote developmental stage: young (< 30 mm), juvenile (30-40 mm), and adult (> 40 mm). Regression lines were calculated separately for each population and sex. Marine Interaction *P* < 0.0001. *P* values for the relationship between SL and tooth number: Premaxilla (males): *P* < 0.0001; Premaxilla (females): *P*=0.121; Dentary (males): *P* < 0.0001; Dentary (females): P=0.553. Freshwater interaction term was nonsignificant; *P* > 0.131. *P* values were determined using an Analysis of Covariance (ANCOVA). Marine: n = 59, freshwater: n = 56.

**Table 3.** Summary of linear regression results for the relationship between tooth number and standard length in marine and freshwater females. Regression estimates, standard errors, t-statistics, and *P*-values are shown for the relationship between tooth number in response to standard length in marine and freshwater females. Slopes represent the rates of tooth addition per mm for each population. Bolded *P*-values highlight the significant relationship between body size and tooth addition in freshwater but not marine females. The significant interaction term (Marine Slope Difference) indicates that the tooth addition in response to increasing standard length in females is population-dependent. Values in parentheses report 95% confidence intervals. The model fit is reported as R^2^ and adjusted R^2^ values. *P*-values were determined using an Analysis of Covariance (ANCOVA). Statistical significance is denoted in bold. Total n=58 (Freshwater n = 29, Marine n = 30).

| *Predictors* | **Premaxilla Total** |  |  |  | **Dentary Total** |  |  |  |
| --- | --- | --- | --- | --- | --- | --- | --- | --- |
|  | *Estimates* | *std. Error* | *Statistic* | *p* | *Estimates* | *std. Error* | *Statistic* | *p* |
| Freshwater Slope | 1.32<br>(0.81 – 1.84) | 0.26 | 5.13 | **<0.0001** | 0.90<br>(0.60 – 1.21) | 0.15 | 6.01 | **<0.0001** |
| Marine Slope | 0.35<br>(-0.10 – 0.80) | 0.22 | 1.57 | 0.1213 | 0.08<br>(-0.18 – 0.34) | 0.13 | 0.60 | 0.5532 |
| Marine Slope Difference (Interaction) | -0.98<br>(-1.66 – -0.29) | 0.34 | -2.87 | **0.0058** | -0.83<br>(-1.22 – -0.43) | 0.20 | -4.17 | **0.0001** |
| Observations | 58 |  |  |  | 58 |  |  |  |
| R<sup>2</sup> / R<sup>2</sup>adjusted | 0.376 / 0.342 |  |  |  | 0.408 / 0.375 |  |  |  |

Given that standard length influenced tooth number in freshwater but not marine females, we were unable to apply a baseline standard length correction to analyze female tooth number differences. However, we identified a crossover point at ∼ 40 mm at which point the oral tooth number in both populations intersected. Thus, we analyzed tooth number differences between females below and above this crossover point (**Tables 4-5**). We found that prior to adulthood (below 40 mm), tooth number in the premaxilla was not significantly different between marine and freshwater females (**Table 4**). However, dentary tooth number was significantly higher in marine populations (**Table 4**). After adulthood (> 40 mm), freshwater females had significantly more premaxillary and dentary teeth (**Figure 11. A-B**; **Table 5**). As development progressed, freshwater females continued gaining teeth and thus surpassed marine females, whose tooth number remained relatively constant.

**Table 4.** Linear model results for population-driven differences in oral tooth number in young and juvenile female sticklebacks. Regression estimates, standard errors, 95% Confidence Intervals (CI), t-statistics, and *P*-values are shown for premaxillary and dentary tooth number in young and juvenile females (< 40 mm) from marine and freshwater populations. Estimates (β) indicate the mean tooth number for freshwater populations and the mean difference in tooth number in the marine population. Values in parentheses report 95% confidence intervals. Bolded *P*-value indicates a significant difference in dentary tooth number between marine and freshwater females. *P*-values were determined using an Analysis of Covariance (ANCOVA). The model fit is reported as R^2^ and adjusted R^2^ values. Total n=33 (Marine n=19, Freshwater n=14).

| *Model Term* | **Premaxilla Total** |  |  |  |  | **Dentary Total** |  |  |  |  |
| --- | --- | --- | --- | --- | --- | --- | --- | --- | --- | --- |
|  | *Estimate*<br>*($\beta$)* | *std.*<br>*Error* | *CI* | *Statistic* | *p* | *Estimate*<br>*($\beta$)* | *std.*<br>*Error* | *CI* | *Statistic* | *p* |
| Freshwater Mean | 35.43 | 2.12 | 31.10 – 39.76 | 16.70 | **<0.001** | 24.14 | 1.07 | 21.97 – 26.32 | 22.62 | **<0.001** |
| Marine (Difference) | 1.52 | 2.80 | -4.19 – 7.22 | 0.54 | 0.591 | 3.38 | 1.41 | 0.51 – 6.25 | 2.41 | **0.022** |
| Observations | 33 |  |  |  |  | 33 |  |  |  |  |
| R<sup>2</sup> / R<sup>2</sup> adjusted | 0.009 / -0.023 |  |  |  |  | 0.157 / 0.130 |  |  |  |  |

**Table 5.** Linear model results for oral tooth number differences in adult marine and freshwater female sticklebacks. Regression estimates, standard errors, 95% Confidence Intervals (CI), t-statistics, and *P*-values are shown for premaxillary and dentary tooth number in adult females (> 40 mm) from marine and freshwater populations. Estimates (β) indicate the mean tooth number in each population and the mean difference in tooth number between populations. Values in parentheses report 95% confidence intervals. Bolded *P*-values indicate the significant difference in premaxillary and dentary tooth number between populations. *P*-values were determined using an Analysis of Covariance (ANCOVA). The model fit i s reported as R^2^ and adjusted R^2^ values. Total n=25 (Marine n=11, Freshwater n=14).

| *Model Term* | **Premaxilla Total** |  |  |  |  | **Dentary Total** |  |  |  |  |
| --- | --- | --- | --- | --- | --- | --- | --- | --- | --- | --- |
|  | *Estimate*<br>*($\beta$)* | *std.*<br>*Error* | *95% CI* | *Statistic* | *p* | *Estimate*<br>*($\beta$)* | *std.*<br>*Error* | *95% CI* | *Statistic* | *p* |
| Freshwater Mean | 51.50 | 2.10 | 47.15 – 55.85 | 24.50 |  | 34.57 | 1.53 | 31.41 – 37.74 | 22.59 |  |
| Marine Mean | 43.5 | 2.37 | 38.5 – 48.4 | 18.33 |  | 29.5 | 1.73 | 26.0 – 33.1 | 17.11 |  |
| Marine (Difference) | -8.05 | 3.17 | -14.60 – -1.49 | -2.54 | **0.018** | -5.03 | 2.31 | -9.80 – -0.25 | -2.18 | **0.046** |
| Observations | 25 |  |  |  |  | 25 |  |  |  |  |
| R<sup>2</sup> / R<sup>2</sup> adjusted | 0.219 / 0.185 |  |  |  |  | 0.171 / 0.135 |  |  |  |  |

Given that standard length influenced tooth number in males from both marine and freshwater populations (**Figure 11**; **Table 6**), we corrected for standard length to determine oral tooth number differences driven by population. Ultimately, we found that freshwater males had significantly more premaxillary teeth than marine males (*P* = 0.0007; **Table 7**). Dentary tooth number was also higher in freshwater males, but was only marginally significant (*P* = 0.068, **Table 7**).

**Table 6.** Summary of linear regression analysis of oral tooth number in response to body size in marine and freshwater male sticklebacks. Regression estimates, standard errors, t-statistics, and *P*-values are shown for the relationship between tooth number in response to standard length in each population. Slopes represent the rates of tooth addition per mm for each population. Bolded *P*-values highlight the significant relationship between body size and tooth addition in each population. Non-significant interaction term (Marine Slope Difference) indicates the non-significant slope difference between populations. Values in parentheses report 95% confidence intervals. The model fit is reported as R^2^ and adjusted R^2^ values. *P*-values were determined using an Analysis of Covariance (ANCOVA). Statistical significance is denoted in bold. Total n=57 (Marine = 29, Freshwater = 28).

| *Predictors* | **Premaxilla Total** |  |  |  | **Dentary Total** |  |  |  |
| --- | --- | --- | --- | --- | --- | --- | --- | --- |
|  | *Estimates* | *std. Error* | *Statistic* | *p* | *Estimates* | *std. Error* | *Statistic* | *p* |
| Freshwater Slope | 1.79<br>(1.48 – 2.11) | 0.16 | 11.50 | **<0.0001** | 1.41<br>(1.16 – 1.65) | 0.12 | 11.36 | **<0.0001** |
| Marine Slope | 1.77<br>(1.37 – 2.17) | 0.20 | 9.0 | **<0.0001** | 1.08<br>(0.76 – 1.40) | 0.16 | 6.88 | **<0.0001** |
| Marine Slope Difference (Interaction) | -0.02<br>(-0.53 – 0.48) | 0.25 | -0.10 | 0.9222 | -0.33<br>(-0.73 – 0.07) | 0.20 | -1.65 | 0.1057 |
| Observations | 57 |  |  |  | 57 |  |  |  |
| R<sup>2</sup> / R<sup>2</sup> adjusted | 0.803 / 0.791 |  |  |  | 0.769 / 0.756 |  |  |  |

**Table 7.** Linear model results for oral tooth number differences between marine and freshwater male sticklebacks. . Regression estimates, standard errors, 95% Confidence Intervals (CI), t-statistics, and *P*-values are shown for total premaxillary and dentary tooth number. Slope indicates the common rate of tooth addition between populations. Marine (Difference) indicates the difference in mean tooth number in marine males relative to freshwater males. Values in parentheses report 95% confidence intervals. Bolded *P*-values indicate the significant relationship between tooth number and body size in both populations, and higher mean premaxillary tooth number in freshwater males. *P*-values were determined using an Analysis of Covariance (ANCOVA). The model fit is reported as R^2^ and adjusted R^2^ values. Total n=57 (Marine=29, Freshwater n=28).

| *Predictors* | **Premaxilla Total** |  |  |  | **Dentary Total** |  |  |  |
| --- | --- | --- | --- | --- | --- | --- | --- | --- |
|  | *Estimates* | *std.*<br>*Error* | *Statistic* | *p* | *Estimates* | *std.*<br>*Error* | *Statistic* | *p* |
| Slope | 1.78<br>(1.54 – 2.03) | 0.12 | 14.72 | **<0.0001** | 1.28<br>(1.08 – 1.48) | 0.10 | 12.97 | **<0.0001** |
| Marine<br>(Difference) | -7.23<br>(-11.26 – -3.19) | 2.01 | -3.59 | **0.0007** | -3.05<br>(-6.33 – 0.24) | 1.64 | -1.86 | 0.0682 |
| Observations | 57 |  |  |  | 57 |  |  |  |
| R<sup>2</sup> / R<sup>2</sup> adjusted | 0.803/ 0.795 |  |  |  | 0.757 / 0.748 |  |  |  |

### Evolved differences in oral tooth number between marine and freshwater populations throughout development

Since sexual dimorphism in oral tooth number differed throughout development, we sought to determine whether evolved differences in tooth number between marine and freshwater populations followed a similar trajectory. Thus, we analyzed oral tooth numbers in marine and freshwater populations from the previously delineated developmental stages: early, late juvenile, and adult stages, analyzing each sex separately.

In females, population did not significantly impact premaxillary tooth number in the young or juvenile stage (**Figure 12**; **Table 8**). However, by adulthood, freshwater females had significantly more premaxillary teeth than marine females (+8.04, *P* = 0.018). Our earlier results showed that below 40 mm, marine females had significantly more dentary teeth than freshwater females (**Table 4**). Looking earlier in development revealed that the elevated dentary tooth number in marine females relative to freshwater females (+6.733, *P* = 0.016) was present at the earliest stage (< 30 mm). The difference in dentary tooth number then became non-significant during the late-juvenile stage, as the slopes of the lines neared each other, after which the developmental trajectory shifted with adult freshwater females surpassing marine females (+5.03, *P* = 0.04); **Figure 12**).

**Figure 12.**
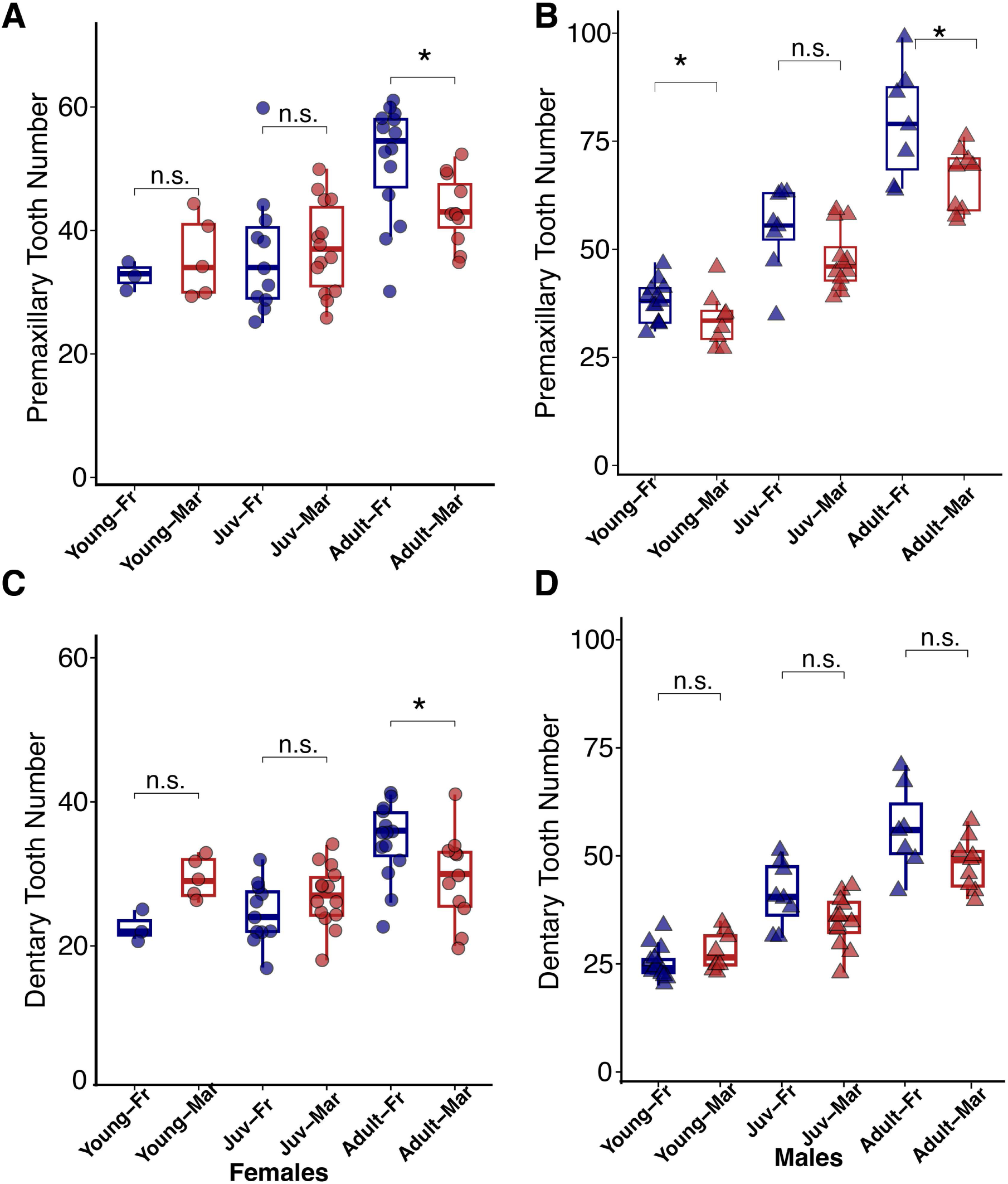
Evolved differences in oral tooth number between marine and freshwater populations throughout development. (A-B) Premaxillary and (C-D) dentary tooth number by population and sex. Mar = marine (red); Fr = freshwater (blue). Females = circles; Males = triangles. Boxes represent 25th-75th percentiles, midlines represent the median, whiskers denote boundaries of the min/max range. Standard length did not significantly influence female tooth number. Raw tooth counts were used for analysis. Standard length significantly influenced tooth number in young and late juvenile males. Size-corrected values were used for the analysis. All *P* values were determined using an Analysis of Covariance (ANCOVA). * *P*<0.05, \*\**P*<0.01, \*\*\**P*<0.001, \*\*\*\**P*<0.0001, n.s. not significant. Female total: n=58. Young n=8 (Fr=3, Mar=5); juvenile n=25 (Fr=11, Mar=14); adult n=25 (Fr=14, Mar=11). Male total n=57. Young n=21 (Fr=13, Mar=8); juvenile n=20 (Fr=8, Mar=12); adult n=16 (Fr=7, Mar=9).

**Table 8.** Linear model analysis of oral tooth number differences in marine and freshwater female sticklebacks across three discrete developmental stages. Regression estimates, standard errors, 95% Confidence Intervals (CI), t-statistics, and *P*-values are shown for premaxillary and dentary tooth number in female sticklebacks from young (<30 mm); juvenile (30-40 mm); and adult (> 40 mm) stages. Estimates (β) represent the mean tooth number and difference in mean tooth number between populations. Values in parentheses report 95% confidence intervals. Bolded *P*-values indicate statistically significant differences in tooth number. *P*-values were determined using an Analysis of Covariance (ANCOVA). The model fit is reported as R^2^ and adjusted R^2^ values. Total n=57 (early= 8; juvenile=25; adult = 25).

| *Model Term* | **Premaxilla Total** |  |  |  |  | **Dentary Total** |  |  |  |  |
| --- | --- | --- | --- | --- | --- | --- | --- | --- | --- | --- |
|  | *Estimate*<br>*($\beta$)* | *std.*<br>*Error* | *95% CI* | *Statistic* | *p* | *Estimate*<br>*($\beta$)* | *std.*<br>*Error* | *95% CI* | *Statistic* | *p* |
| **Young (< 30mm)** |  |  |  |  |  |  |  |  |  |  |
| Freshwater Mean | 32.67 | 3.25 | 24.72 – 40.61 | 10.06 |  | 22.67 | 1.60 | 18.76 – 26.57 | 14.20 |  |
| Marine Mean | 35.6 | 2.52 | 29.4 – 41.8 | 14.15 |  | 29.4 | 1.24 | 26.4 – 32.4 | 23.78 |  |
| Marine (Difference) | 2.93 | 4.11 | -7.12 – 12.99 | 0.71 | 0.502 | 6.73 | 2.02 | 1.79 – 11.67 | 3.33 | **0.016** |
| Observations | 8 |  |  |  |  | 8 |  |  |  |  |
| R<sup>2</sup> / R<sup>2</sup> adjusted | 0.078 / -0.075 |  |  |  |  | 0.650 / 0.591 |  |  |  |  |
| **Juvenile (30-40mm)** |  |  |  |  |  |  |  |  |  |  |
| Freshwater Mean | 36.18 | 2.61 | 30.78 – 41.58 | 13.86 |  | 24.55 | 1.28 | 21.89 – 27.20 | 19.13 |  |
| Marine Mean | 37.4 | 2.31 | 32.6 – 42.2 | 16.18 |  | 26.9 | 1.14 | 24.5 – 29.2 | 23.62 |  |
| Marine (Difference) | 1.25 | 3.49 | -5.97 – 8.46 | 0.36 | 0.724 | 2.31 | 1.71 | 1.23 – 5.86 | 1.35 | 0.191 |
| Observations | 25 |  |  |  |  | 25 |  |  |  |  |
| R<sup>2</sup> / R<sup>2</sup> adjusted | 0.006 / -0.038 |  |  |  |  | 0.073 / 0.033 |  |  |  |  |
| **Adult (> 40mm)** |  |  |  |  |  |  |  |  |  |  |
| Freshwater Mean | 51.50 | 2.10 | 47.15 – 55.85 | 24.50 |  | 34.57 | 1.54 | 31.41 – 37.74 | 22.59 |  |
| Marine Mean | 43.5 | 2.37 | 38.5 – 48.4 | 18.33 |  | 29.5 | 1.73 | 26.0 – 33.1 | 17.11 |  |
| Marine (Difference) | 8.05 | 3.17 | -14.60 – 1.49 | 2.54 | **0.018** | 5.03 | 2.31 | 9.80 – 0.25 | 2.18 | **0.046** |
| Observations | 25 |  |  |  |  | 25 |  |  |  |  |
| R<sup>2</sup> / R<sup>2</sup> adjusted | 0.219 / 0.185 |  |  |  |  | 0.171 / 0.135 |  |  |  |  |

The evolved differences in oral tooth number between freshwater and marine males followed a different trajectory, with both young and adult freshwater males having significantly more premaxillary teeth than marine males (young: +6.10, *P* = 0.0049; adult: +13.36, *P* = 0.021; **Figure 12**; **Table 9**), while there was no significant difference in the late-juvenile stage (*P* = 0.71). Unlike in females, population did not significantly influence dentary tooth number in males during any stage (**Figure 12**; **Table 9**).

**Table 9.** Linear model analysis of oral tooth number differences in marine and freshwater male sticklebacks across three discrete developmental stages. Regression estimates, standard errors, 95% Confidence Intervals (CI), t-statistics, and *P*-values are shown for premaxillary and dentary tooth number in marine and freshwater male sticklebacks from young (<30 mm); juvenile (30-40 mm); and adult (> 40 mm) stages. Estimates (β) represent mean tooth number in each population and the difference in tooth number between populations at a given stage. Values in parentheses report 95% confidence intervals. Bolded *P*-values indicate statistically significant differences in tooth number between populations. The model fit is reported as R^2^ and adjusted R^2^ values. *P*-values were determined using an Analysis of Covariance (ANCOVA). n=57 (early= 21; juvenile=20; adult n= 16).

| *Model Term* | **Premaxilla Total** |  |  |  |  | **Dentary Total** |  |  |  |  |
| --- | --- | --- | --- | --- | --- | --- | --- | --- | --- | --- |
|  | *Estimate*<br>*($\beta$)* | *std.*<br>*Error* | *95% CI* | *Statistic* | *p* | *Estimate*<br>*($\beta$)* | *std.*<br>*Error* | *95% CI* | *Statistic* | *p* |
| **Young (< 30mm)** |  |  |  |  |  |  |  |  |  |  |
| Freshwater Mean | 38.6 | 1.15 | 36.2 – 41 | 33.76 |  | 25.8 | 0.89 | 24.0 – 27.7 | 29.09 |  |
| Marine Mean | 32.5 | 1.47 | 29.4 – 35.6 | 22.04 |  | 27.0 | 1.14 | 24.6 – 29.4 | 23.66 |  |
| Marine<br>(Difference) | -6.11 | 1.90 | -10.10 –<br>-2.11 | -3.21 | **0.005** | 1.22 | 1.47 | -1.88 – 4.31 | 0.82 | 0.420 |
| Observations | 21 |  |  |  |  | 21 |  |  |  |  |
| R<sup>2</sup> / R<sup>2</sup> adjusted | 0.516 / 0.462 |  |  |  |  | 0.499 / 0.443 |  |  |  |  |
| **Juvenile (30-40mm)** |  |  |  |  |  |  |  |  |  |  |
| Freshwater Mean | 49.7 | 2.40 | 44.6 – 54.7 | -20.709 |  | 39.5 | 2.66 | 33.9 – 45.1 | 14.88 |  |
| Marine Mean | 50.9 | 1.88 | 46.9 – 54.9 | -27.09 |  | 35.9 | 2.08 | 31.5 – 40.3 | 17.26 |  |
| Marine<br>(Difference) | 1.25 | 3.33 | -5.78 – 8.28 | 0.37 | 0.713 | -3.60 | 3.69 | -11.38 –<br>4.19 | -0.97 | 0.343 |
| Observations | 20 |  |  |  |  | 20 |  |  |  |  |
| R<sup>2</sup> / R<sup>2</sup> adjusted | 0.582 / 0.532 |  |  |  |  | 0.243 / 0.154 |  |  |  |  |
| **Adult (> 40mm)** |  |  |  |  |  |  |  |  |  |  |
| Freshwater Mean | 79.14 | 3.86 | 70.87 –<br>87.41 | 20.53 |  | 56.29 | 3.03 | 49.79 –<br>62.78 | 18.58 |  |
| Marine Mean | 65.8 | 3.40 | 58.5 – 73.1 | 19.35 |  | 48.2 | 2.67 | 42.5 – 54.0 | 18.05 |  |
| Marine<br>(Difference) | -13.37 | 5.14 | -24.39 –<br>-2.34 | -2.60 | **0.021** | -8.06 | 4.04 | -16.72 –<br>0.60 | -2.00 | 0.066 |
| Observations | 16 |  |  |  |  | 16 |  |  |  |  |
| R<sup>2</sup> / R<sup>2</sup> adjusted | 0.326 / 0.277 |  |  |  |  | 0.222 / 0.166 |  |  |  |  |

### Differences in oral tooth number of marine adults by seasonal lighting conditions

Given the role of oral teeth in nest-building and intersex fighting in male sticklebacks during breeding season^36,47–49^, we compared tooth number in adult (> 40 mm) marine fish from “winter” (8 hours of light per day) and “summer” (16 hours of light per day) families. We discovered that in both sexes, fish from summer conditions had significantly more premaxillary and dentary teeth than those from winter conditions (**Figure 13**; **Tables 10-11**). While summer conditions led to increased tooth number in both sexes, males responded more strongly to this increase in numbers of light per day than females (*P* = 0.023; S: +37.41; W: +22.32). Ultimately, this led to greater sexual dimorphism present in the premaxilla of summer than winter fish (**Figure 13**; **Table 12**). In the dentary, tooth number was significantly higher in summer than winter (F: +9.41, *P* < 0.001; M: +11.69, *P* = 8.61e3); however, the increase in dentary tooth number in summer fish was uniform across both sexes (*P* = 0.59; S: +20.95; W: +18.68). We also observed greater variation in oral tooth number in males in summer conditions than in winter conditions, perhaps correlating with male’s secondary sexual characteristics.

**Figure 13.**
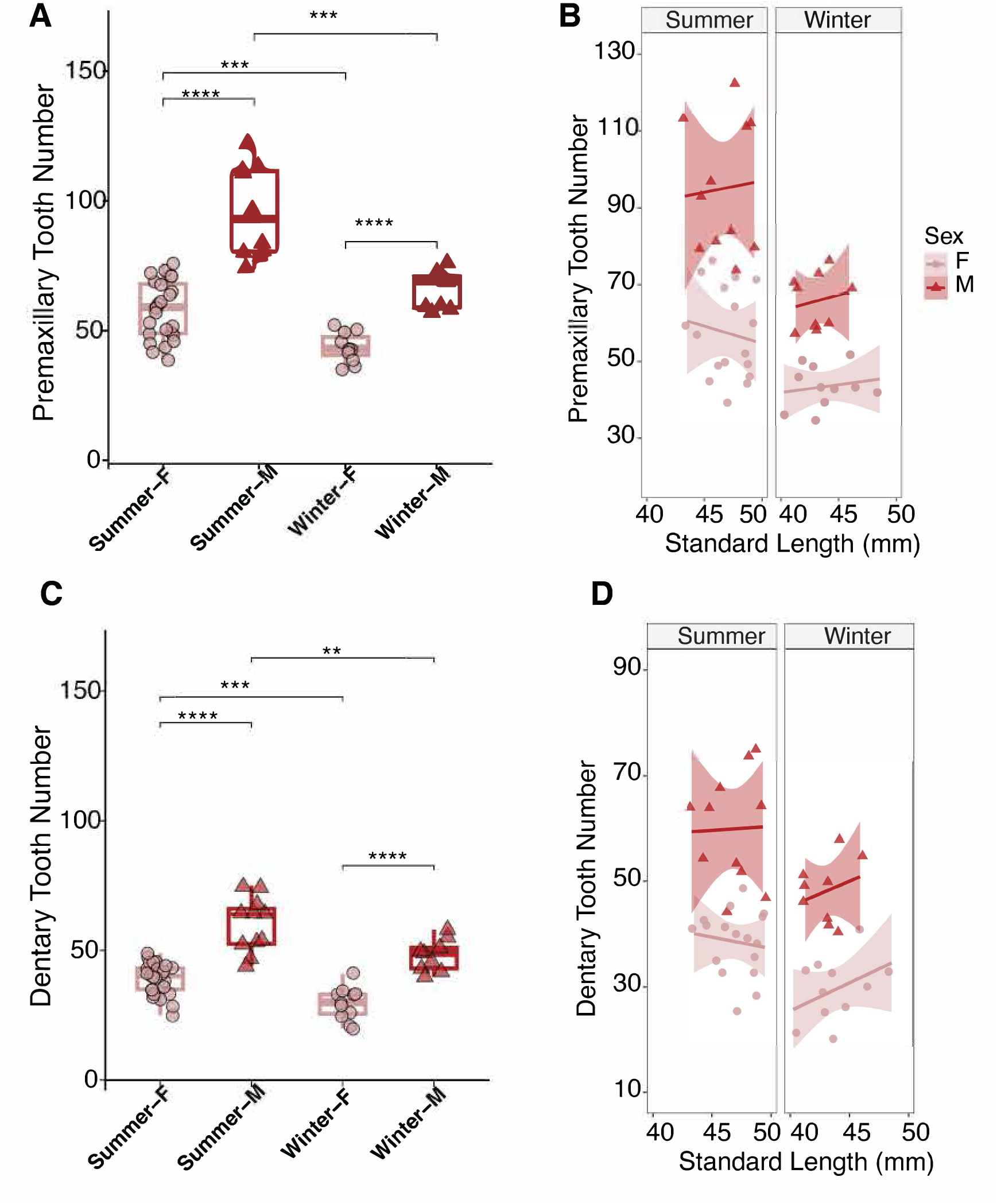
Increased oral tooth number in marine fish raised in summer conditions. (A-B) Premaxillary and (C-D) dentary tooth counts in adult marine fish from “summer” (16 hours of light per day) and “winter” (8 hours of light per day) raised families. (A,C) Boxplots indicate raw tooth counts from the 25th-75th percentiles, midlines represent the median, and whiskers denote boundaries of the min/max range. *P* values were determined using an Analysis of Covariance (ANCOVA). * *P*<0.05, \*\**P*<0.01, \*\*\**P*<0.001, \*\*\*\**P*<0.0001, n.s. not significant. Winter: n = 36 (F=25, M=11); Summer n = 20 (F=11, M=9). (B, D) Linear regression of oral tooth counts relative to standard length (n.s.) in adult marine fish. Shaded regions indicate 95% confidence intervals. Points represent individual fish (triangles/dark red = males; circles/light = females). F = Females; M = Males; S = Summer; W = Winter.

**Table 10.** Linear model results for the effect of seasonality on oral tooth number in female marine sticklebacks. . Regression estimates, standard errors, 95% Confidence Intervals (CI), t-statistics, and *P*-values are shown for premaxillary and dentary tooth number in adult marine females (> 40 mm) from "summer" (16 hours of light per day) and "winter" (8 hours of light per day) raised families. Estimates (β) indicate the mean tooth number in fish from each season and mean difference in tooth number seasons. Values i n parentheses report 95% confidence intervals. Bolded *P*-values indicate the significant increase in tooth number in "summer" raised fish. *P*-values were determined using an Analysis of Covariance (ANCOVA). The model fit is reported as R^2^ and adjusted R^2^ values. Total n=36 (Winter=11, Summer=25).

| *Model Term* | **Premaxilla Total** |  |  |  |  | **Dentary Total** |  |  |  |  |
| --- | --- | --- | --- | --- | --- | --- | --- | --- | --- | --- |
|  | *Estimate*<br>*($\beta$)* | *std.*<br>*Error* | *95% CI* | *Statistic* | *p* | *Estimate*<br>*($\beta$)* | *std.*<br>*Error* | *95% CI* | *Statistic* | *p* |
| Summer Mean | 57.68 | 1.95 | 53.73 –<br>61.63 | 29.64 |  | 38.96 | 1.24 | 36.44 –<br>41.48 | 31.43 |  |
| Winter Mean | 43.5 | 2.93 | 37.5 –<br>49.4 | 14.81 |  | 29.5 | 1.87 | 25.7 –<br>33.3 | 15.81 |  |
| Winter<br>(Difference) | -14.23 | 3.52 | -21.38 –<br>-7.07 | -4.04 | **<0.001** | -9.41 | 2.24 | -13.97 –<br>-4.86 | -4.20 | **<0.001** |
| Observations | 36 |  |  |  |  | 36 |  |  |  |  |
| R<sup>2</sup> / R<sup>2</sup> adjusted | 0.324 / 0.305 |  |  |  |  | 0.341 / 0.322 |  |  |  |  |

**Table 11.** Linear model results for the effect of seasonality on oral tooth number in male marine sticklebacks. Regression estimates, standard errors, 95% Confidence Intervals (CI), t-statistics, and *P*-values are shown for premaxillary and dentary tooth number in adult marine males (> 40 mm) from "summer" (16 hours of light per day) and "winter" (8 hours of light per day) raised families. Estimates (β) indicate the mean tooth number in males from each season and difference in tooth number between summer and winter populations. Values in parentheses report 95% confidence intervals. Bolded *P*-values indicate the significant increase in oral tooth number in "summer" populations. *P*-values were determined using an Analysis of Covariance (ANCOVA). The model fit is reported as R^2^ and adjusted R^2^ values. Total n=20 (Winter n=9, Summer n=11).

| *Model Term* | **Premaxilla Total** |  |  |  |  | **Dentary Total** |  |  |  |  |
| --- | --- | --- | --- | --- | --- | --- | --- | --- | --- | --- |
|  | *Estimate*<br>*($\beta$)* | *std.*<br>*Error* | *95% CI* | *Statistic* | *p* | *Estimate*<br>*($\beta$)* | *std.*<br>*Error* | *95% CI* | *Statistic* | *p* |
| Summer Mean | 95.09 | 4.06 | 86.56 –<br>103.62 | 23.41 |  | 59.91 | 2.66 | 54.33 –<br>65.49 | 22.54 |  |
| Winter Mean | 65.8 | 4.49 | 56.3 –<br>75.2 | 14.65 |  | 48.2 | 2.94 | 42.0 –<br>54.4 | 16.41 |  |
| Winter<br>(Difference) | -29.31 | 6.05 | -42.03 –<br>-16.59 | -4.84 | **<0.001** | -11.69 | 3.96 | -20.01 –<br>-3.36 | -2.95 | **0.009** |
| Observations | 20 |  |  |  | 20 |  |  |  |  |  |
| R<sup>2</sup> / R<sup>2</sup> adjusted | 0.566 / 0.541 |  |  |  | 0.326 / 0.288 |  |  |  |  |  |

**Table 12.** Summary of linear regression analysis of sexual dimorphism in oral tooth number in "summer" marine sticklebacks. Regression estimates, standard errors, t-statistics, 95% Confidence Intervals (CI), and *P*-values are shown for tooth number differences between "summer" (16 hours of light per day) males and females. Estimates (β) represent mean tooth number in each sex and the difference in tooth number in males. Bolded *P*-values highlight significant differences in tooth number. The model fit is reported as R^2^ and adjusted R^2^ values. *P*-values were determined using an Analysis of Covariance (ANCOVA). Total n = 36 (Females n=25, Males n=11).

| *Model Term* | **Premaxilla Total** |  |  |  |  | **Dentary Total** |  |  |  |  |
| --- | --- | --- | --- | --- | --- | --- | --- | --- | --- | --- |
|  | *Estimate*<br>*($\beta$)* | *std.*<br>*Error* | *95% CI* | *Statistic* | *p* | *Estimate*<br>*($\beta$)* | *std.*<br>*Error* | *95% CI* | *Statistic* | *p* |
| Female Mean | 57.68 | 2.60 | 52.39 –<br>62.97 | 22.16 |  | 38.96 | 1.54 | 35.82 –<br>42.10 | 25.22 |  |
| Male Mean | 95.1 | 3.92 | 87.1 –<br>103 | 24.23 |  | 59.99 | 2.33 | 55.2 –<br>64.6 | 25.73 |  |
| Male<br>Difference | +37.41 | 4.71 | 27.84 –<br>46.98 | 7.94 | **<0.001** | +20.95 | 2.79 | 15.27 –<br>26.63 | 7.50 | **<0.001** |
| Observations |  | 36 |  |  |  |  | 36 |  |  |  |
| R<sup>2</sup> / R<sup>2</sup> adjusted |  |  | 0.650 / 0.640 |  |  |  |  | 0.623 / 0.612 |  |  |

## DISCUSSION

### Spatial and temporal development of oral teeth

Tooth patterning and morphology is highly variable within teleosts, with certain groups having both oral and pharyngeal teeth, while others possess only pharyngeal teeth^50^. We previously described significant evolved tooth gain in the pharyngeal jaws of freshwater stickleback populations^37,38^, raising questions about whether the oral jaws also evolve tooth gain in freshwater sticklebacks. Furthermore, since sticklebacks have both an oral and pharyngeal jaw, determining the early spatiotemporal patterning of primary tooth formation in the oral jaw allowed us to assess the degree of similarity between the two jaws. While both pharyngeal and oral teeth replace throughout the lifespan, the timing of the first replacement event on the oral jaw, as well as the spatiotemporal sequence of primary oral tooth formation was not known. We observed that tooth formation on the oral jaw occurs more slowly than on the pharyngeal jaw. Unlike pharyngeal teeth, which form prior to hatching, oral teeth do not form until 1-3 days after hatching. Tooth maturation then progresses more slowly. While in the pharyngeal jaw, the pioneer tooth on the Ventral Tooth Plate (VTP), Dorsal Tooth Plate 2 (DTP2), and DTP1 calcify by 8 and 12 dpf, respectively, in the oral jaw, the pioneer tooth does not calcify on the premaxilla and dentary until 13-14 dpf^15^ 20. Similarly, in the pharyngeal jaw, by 20 dpf, VTP and DTP2 possess between 14-20 teeth and DTP1 possesses 8 teeth, while in both the premaxilla and dentary, only 6-8 teeth are observed at this stage^15^.

In both the upper and lower jaw, a new tooth formed roughly every two days, with most fish having between five to six teeth on each jaw by 12 days after hatching (28 dpf). Between 20-28 dpf the rate of tooth formation slowed, with most fish still having between six to seven teeth by the end of this time period. However, the maturation stage of the first four teeth progressed, with a seventh tooth only forming after the first five became ankylosed. This timing may relate to the different replacement mechanisms that have been identified between the oral and pharyngeal jaws. While in the pharyngeal jaw, replacement teeth dislodge the tooth they are replacing, in the oral jaw, replacement teeth have been suggested to form in a one-to-one fashion^25^. Given the different functions that the pharyngeal and oral dentition serve, the former being involved in prey mastication and the latter being evolved in prey capture and nest building, there is the potential for evolution to act differently in these dentitions^18–21^.

Two primary hypotheses have been proposed regarding the mechanism by which teeth form. In the Zahnreihen model, the first tooth acts as a signaling center, inducing the formation of subsequent teeth in a wave-like fashion due to the secretion of activating factors^51–54^. A second mechanism for the establishment of the tooth field is the zone of inhibition or reaction diffusion model, wherein the interplay between activating and inhibiting signaling factors controls where a tooth forms^55–60^. It is also possible that the mechanism of tooth formation differs between primary and replacement teeth^61^.

We observed consistent spatial trends in primary tooth development in both the premaxilla and dentary. In both jaws, the distance between the pioneer and second tooth was significantly shorter than the distance between the pioneer and the third tooth (**Figure S6**). Therefore, it is possible that inhibitory signals from both the pioneer and second tooth result in the third tooth forming at a greater distance than would be expected if only one tooth emerged before. The close time window in which the first three teeth form also increases the likelihood of residual inhibitory factors from when the first two teeth formed.

Even at the start of oral tooth formation, we observed evidence of uncoupling between the premaxilla and dentary. In several instances, we observed a tooth initiating on only one of the jaws. These observations are in line with previous studies that have shown that in the bichir *Polypterus senegalus*, primary teeth form on the dentary before the premaxilla^62^. The jaw decoupling we observed persisted throughout development, with nearly 45-55% of fish between 8-28 dpf exhibiting decoupling in premaxillary and dentary tooth number. Interestingly, we observed a strong directional bias, with 84.8-89.2% of instances of decoupling occurring when the dentary possessed more teeth than the premaxilla in marine and freshwater populations, respectively. Perhaps as a result of the increased rate of tooth formation on the dentary, we also observed several instances of tooth shedding and replacement in the dentary, when no such instances had occurred in the premaxilla. This decoupling is similar to what has been observed in elasmobranchs, where the rates of replacement may differ between upper and lower jaws^63^.

Further, tooth positioning on the dentary was more variable than on the premaxilla. In the premaxilla the position of the second tooth was highly stereotypical, occurring medial to the pioneer tooth in most instances, whereas on the dentary, the second tooth occupied the medial and lateral positions with roughly equal frequency. Similarly, the fourth tooth on the dentary occupied one of three positions, while on the premaxilla, the fourth tooth primarily erupted lateral to the third tooth. Instances of uncoupling between the two jaws have also been observed in other teleosts^9,64,65^.

Overall, the spatiotemporal sequence of early oral tooth formation was similar between the ancestral marine and derived freshwater populations. However, we did observe increased variability in the dentary with respect to the second tooth, which most often formed laterally to the pioneer in freshwater, and medial to the pioneer in marine fish.

### Sexual dimorphism arises late in development

Expectedly, neither sex nor population influenced tooth number in fish below 30 mm. However, by the juvenile stage (30-40 mm), both marine and freshwater males had significantly more oral teeth than females. Given that this difference in tooth number arises near the timing of gonad differentiation, it is possible that changes in hormone levels may contribute to the sexual dimorphism we observed^66,67^. This late emerging dimorphism is consistent with what has been observed in other teleost species where differences in pelvic girdle, body coloring, and overall body size increase first emerge near stages of gonad differentiation^31,32,68^. Interestingly, significant differences in body size were not found in the juvenile stage, suggesting that the increase in tooth number is not solely a result of body size increases.

We also observed sexual dimorphism in the relationship between tooth number and body size between the ancestral marine and derived freshwater populations. While in freshwater fish, both sexes exhibited increases in tooth number with increasing body size, in the marine population, only males displayed a significant relationship between body size and tooth number. Given the selective pressures imposed on the derived freshwater population as a result of rapid and repeated colonization of new environments, these selective pressures might override the pressures of sexual selection^69^. Further, the fact that these selective pressures affect both sexes uniformly, we would expect to see females become more male-like over time. This divergence model proposes that the degree of sexual dimorphism should decrease in the derived population relative to the ancestral population^27,30^. Indeed, recent work exploring the sexually dimorphic traits of body size, pelvic girdle, and jaw length found that sexual dimorphism was significantly greater in the marine population relative to the freshwater population^34,35^. However, the sexual dimorphism in oral tooth number appeared even more significant in freshwater fish than marine fish, not supporting this divergence model.

### Evolved tooth gain in the oral jaw

Similar to the pharyngeal jaw, we also observed increases in oral tooth number in adults from the derived freshwater population relative to the ancestral marine population^37,38^. These differences were also influenced by sex and provided support for potential uncoupling between the upper and lower oral jaws. Whereas adult freshwater females had significantly more premaxillary and dentary teeth than marine females, in adult males, tooth number was only significantly different in the premaxilla. This finding suggests that the mechanisms underlying tooth formation in the premaxilla and dentary may differ.

### Effects of lighting upon tooth number

We found that irrespective of body size, “summer” (16 hours of light per day) or “winter” (8 hours of light per day) rearing conditions significantly influenced tooth number in marine fish of both sexes. It is known that selection on dimorphic traits may fluctuate between breeding season and other times of the year^70^. Therefore, it is possible that shifts in hormone levels contribute in part to the changes observed in tooth number in the summer months. While in the dentary, the season-driven increase in tooth number was similar in both sexes, in the premaxilla, the season-driven increase in tooth number was significantly greater in males. During breeding season, males exhibit unique morphological and behavioral traits including changes in nuptial coloration, nest building, and courtship rituals^47,71^.Therefore, it is possible that these features result in season influencing premaxillary tooth number differently in males^66,72,73^.

### Summary

Altogether, our findings that the stickleback oral jaw, like the pharyngeal jaw, evolves increases in tooth number in derived freshwater fish relative to ancestral marine fish suggests either shared selective pressures to increase tooth numbers in both jaws and/or developmental constraints that make the two dentitions covary. Further, our findings of significant sexual dimorphism and population differences in oral tooth number in lab-reared fish fed the same diet suggest that both dimorphism and population differences are hard-wired and genetically encoded. Whether the same genes underlie evolved tooth gain in both the oral and pharyngeal jaw, and whether during development these two sets of teeth are regulated similarly, can now be addressed in future genetic studies.

## MATERIALS AND METHODS

### Stickleback fish husbandry and sex genotyping

For all experiments, stickleback *(Gasterosteus aculeatus*) larvae and adults were lab-reared in brackish water at 18℃ (3.5g/l Instant Ocean salt, 0.217 ml/1 10% sodium bicarbonate). For the early oral tooth time course, artificially fertilized clutches were raised in 150 mm x 15 mm petri dishes. Six marine and seven freshwater clutches were artificially fertilized, and a total of 91 marine and 94 freshwater larvae were sampled. Freshwater fish were from the Cerrito Creek (El Cerrito, California, USA) population and marine fish were from the Rabbit Slough (Alaska, USA) population. Fish were raised in both “winter” (8-hrs of light) and “summer” (16-hrs of light) conditions. Beginning three days after hatching, larvae were fed live brine shrimp daily. Adult fish (over ∼20 mm standard length (SL)) raised in winter were fed live brine shrimp and frozen bloodworms, and adult fish (over ∼40 mm SL) raised in summer were fed frozen bloodworms, frozen daphnia, and frozen mysis shrimp. DNA was extracted from the caudal fins of each fish and sex-genotyping was performed using PCR amplification with sex-specific forward (CATATTGCTGCTTGTGTGGAAG) and reverse (GATCCTCCTCGTTCCTACAG) primers, resulting in fragment sizes of 186 bp and 229 bp from X and Y chromosomes, respectively^46^. All experiments were performed with the approval of the Institutional Animal Care and Use Committee of the University of California, Berkeley (AUP-2015-01-7117-3).

### Skeletal staining and visualization

For skeletal staining, fish larvae and adults were fixed in 10% neutral buffered formalin for 4 hours at room temperature. Following fixation, fish were rinsed in tap water for 2 to 24 hours at room temperature. Fish were transferred to 0.008% Alizarin red in 1% potassium hydroxide (KOH) for 1-7 days for skeletal staining. Fish were washed in tap water for a day and then cleared in 1% KOH for a day. Adult fish were further cleared in 50% glycerol in 0.25% KOH for a day. Stepwise washes in 25% glycerol in 1% KOH, 50% glycerol in 1% KOH, and 75% glycerol in 1% KOH were performed for 15 minutes each. Oral jaws from the fish were then dissected and mounted in 100% glycerol, as previously described^15,39^. A Fisher Scientific Traceable caliper #4 was used to measure the standard length of each fish, from snout to caudal peduncle. Premaxilla and dentary tooth number were each visualized using a DM2500 Leica microscope under both a TX2 red filter and/or under brightfield illumination. Fluorescent images were taken on a Leica DM2500 using the TX2 filter, under 10X magnification. Premaxilla and dentary teeth were counted, and statistical analyses were performed in R.

### Early temporal sequence determination

To determine whether early oral tooth development follows a stereotypical sequence similar to pharyngeal tooth development^15^, we conducted a dense developmental time course of wild-type marine and freshwater stickleback larvae. Following fixation and Alizarin staining, the oral jaws were dissected and mounted onto bridged glass coverslips. We determined the primary tooth sequence of the oral jaw by imaging both the premaxilla (upper jaw) and dentary (lower jaw). The left side of each fish was used for the early sequence determination, and the position of each tooth was scored in the order in which it emerged. Tooth sequence was determined through: (1) direct observation of a tooth emerging at a particular time point, as well as (2) the degree of tooth maturation (ankylosis). If the Alizarin staining was contiguous from the base of the tooth to the jawbone, it was scored as ankylosed. When the base of the tooth was not fully attached to the bone, it was scored as a developing newly mineralized tooth. The consensus sequence was determined separately for the marine and freshwater populations. If the majority of fish from a particular time point displayed a tooth at that given position, it was included in the final sequence.

## ACKNOWLEDGEMENTS

We thank Tyler Square, Brynn Brady, Anthony Vo, and the Miller lab for helpful suggestions on this work. This work was supported by National Institutes of Health grant (DE021475 to C.T.M.) and a Ford Foundation Fellowship (to A.M.).

