## Supplementary Materials for "Development, dimorphism, and divergence of the oral dentition in threespine sticklebacks (*Gasterosteus aculeatus*)"

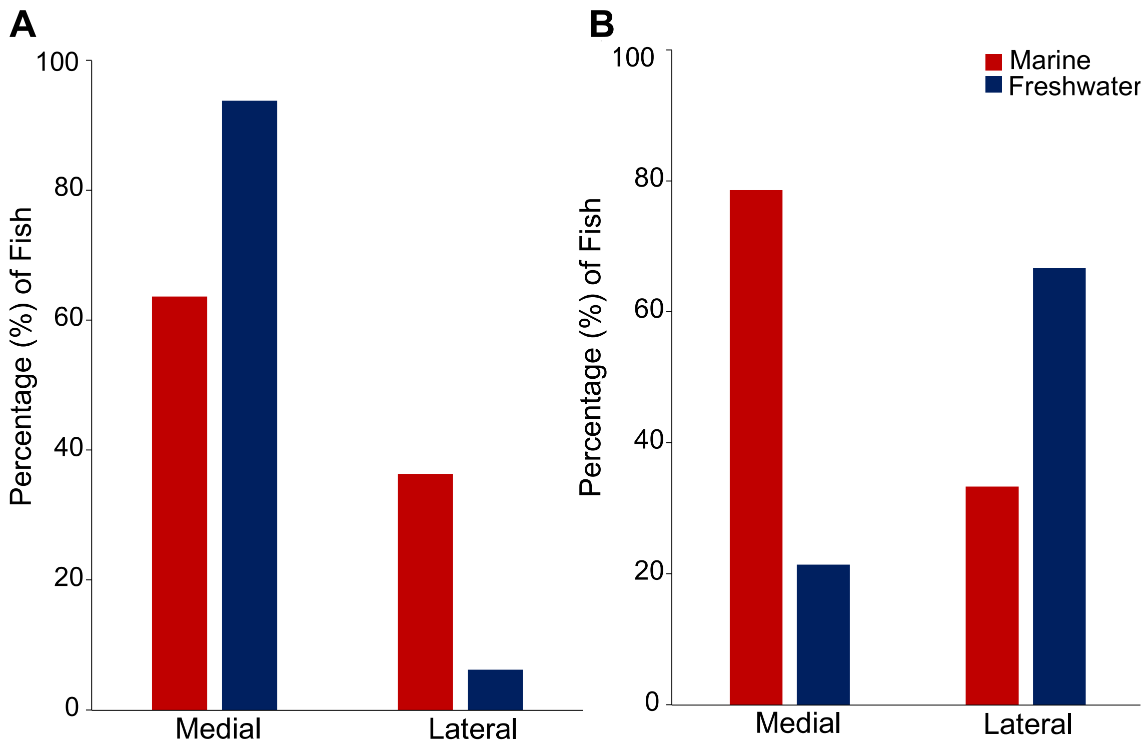


**Figure S1. Positioning of the second tooth relative to the pioneer is consistent between populations in the premaxilla, but exhibits positional variability in the dentary.** (A) Percentage of fish where the second tooth emerged medial (left) or lateral (right) relative to the pioneer on the premaxilla. Marine n=11; Freshwater n=16. (B) Percentage of fish where the second tooth on the dentary emerged medial or lateral to the pioneer. Marine n=15; Freshwater n = 15.

### Figure S2.

| | | | | Marine Premaxilla | | | | | | | | Key |
|---|---|---|---|---|---|---|---|---|---|---|---|---|
| Time Course | Fish ID | Reference | dpf | 1 | 2 | 3 | 4 | 5 | 6 | 7 | 7b | Developing Germ |
| TC5 | 9-1 | 1 | 9 | | | | | | | | | Ankylosed Tooth |
| TC5 | 9-2 | 2 | 9 | | | | | | | | | Absent |
| TC5 | 9-3 | 3 | 9 | | | | | | | | | |
| TC5 | 9-4 | 4 | 9 | | | | | | | | | |
| TC5 | 10-1 | 5 | 10 | | | | | | | | | |
| TC5 | 10-2 | 6 | 10 | | | | | | | | | |
| TC5 | 10-3 | 7 | 10 | | | | | | | | | |
| TC5 | 10-4 | 8 | 10 | | | | | | | | | |
| TC3 | 11-1 | 9 | 11 | | | | | | | | | |
| TC3 | 11-3 | 10 | 11 | | | | | | | | | |
| TC4 | 10-2 | 11 | 11 | | | | | | | | | |
| TC4 | 10-3 | 12 | 11 | | | | | | | | | |
| TC3 | 12-1 | 13 | 12 | | | | | | | | | |
| TC3 | 12-2 | 14 | 12 | | | | | | | | | |
| TC4 | 11-1 | 15 | 12 | | | | | | | | | |
| TC6 | 12-1 | 16 | 12 | | | | | | | | | |
| TC6 | 12-2 | 17 | 12 | | | | | | | | | |
| TC6 | 12-3 | 18 | 12 | | | | | | | | | |
| TC6 | 12-4 | 19 | 12 | | | | | | | | | |
| TC6 | 12-5 | 20 | 12 | | | | | | | | | |
| TC6 | 12-6 | 21 | 12 | | | | | | | | | |
| TC6 | 12-8 | 22 | 12 | | | | | | | | | |
| TC6 | 12-10 | 23 | 12 | | | | | | | | | |
| TC6 | 12-11 | 24 | 12 | | | | | | | | | |
| TC3 | 13-1 | 25 | 13 | | | | | | | | | |
| TC4 | 12-2 | 26 | 13 | | | | | | | | | |
| TC4 | 12-3 | 27 | 13 | | | | | | | | | |
| TC4 | 12-4 | 28 | 13 | | | | | | | | | |
| TC5 | 13-1 | 29 | 13 | | | | | | | | | |
| TC5 | 13-2 | 30 | 13 | | | | | | | | | |
| TC5 | 13-3 | 31 | 13 | | | | | | | | | |
| TC6 | 13-2 | 32 | 13 | | | | | | | | | |
| TC6 | 13-3 | 33 | 13 | | | | | | | | | |
| TC6 | 13-9 | 34 | 13 | | | | | | | | | |
| TC6 | 13-10 | 35 | 13 | | | | | | | | | |
| TC2 | 16-1 | 36 | 14 | | | | | | | | | |
| TC2 | 16-2 | 37 | 14 | | | | | | | | | |
| TC2 | 16-3 | 38 | 14 | | | | | | | | | |
| TC4 | 13-2 | 39 | 14 | | | | | | | | | |
| TC3 | 15-3 | 40 | 15 | | | | | | | | | |
| TC4 | 14-1 | 41 | 15 | | | | | | | | | |
| TC4 | 14-2 | 42 | 15 | | | | | | | | | |
| TC4 | 14-3 | 43 | 15 | | | | | | | | | |
| TC2 | 18-3 | 44 | 16 | | | | | | | | | |
| TC4 | 15-2 | 45 | 16 | | | | | | | | | |
| TC3 | 17-1 | 46 | 17 | | | | | | | | | |
| TC3 | 17-2 | 47 | 17 | | | | | | | | | |
| TC5 | 18-1 | 48 | 18 | | | | | | | | | |
| TC5 | 18-2 | 49 | 18 | | | | | | | | | |
| TC5 | 18-3 | 50 | 18 | | | | | | | | | |
| TC5 | 18-4 | 51 | 18 | | | | | | | | | |
| TC3 | 19-1 | 52 | 19 | | | | | | | | | |
| TC5 | 19-1 | 53 | 19 | | | | | | | | | |
| TC5 | 19-2 | 54 | 19 | | | | | | | | | |
| TC5 | 19-3 | 55 | 19 | | | | | | | | | |
| TC5 | 19-4 | 56 | 19 | | | | | | | | | |
| TC2 | 22-2 | 57 | 20 | | | | | | | | | |
| TC5 | 20-1 | 58 | 20 | | | | | | | | | |
| TC5 | 20-2 | 59 | 20 | | | | | | | | | |
| TC5 | 20-3 | 60 | 20 | | | | | | | | | |
| TC5 | 20-4 | 61 | 20 | | | | | | | | | |
| TC3 | 21-1 | 62 | 21 | | | | | | | | | |
| TC5 | 25-1 | 63 | 25 | | | | | | | | | |
| TC5 | 25-2 | 64 | 25 | | | | | | | | | |
| TC5 | 25-3 | 65 | 25 | | | | | | | | | |
| TC5 | 25-4 | 66 | 25 | | | | | | | | | |
| TC5 | 26-1 | 67 | 26 | | | | | | | | | |
| TC5 | 26-2 | 68 | 26 | | | | | | | | | |
| TC5 | 26-3 | 69 | 26 | | | | | | | | | |
| TC5 | 26-4 | 70 | 26 | | | | | | | | | |
| TC5 | 27-1 | 71 | 27 | | | | | | | | | |
| TC5 | 27-2 | 72 | 27 | | | | | | | | | |
| TC5 | 27-4 | 73 | 27 | | | | | | | | | |
| TC5 | 28-1 | 74 | 28 | | | | | | | | | |
| TC5 | 28-3 | 75 | 28 | | | | | | | | | |
| TC5 | 28-4 | 76 | 28 | | | | | | | | | |

**Figure S2. Scoring of the primary premaxillary tooth sequence in marine larvae.** Numbers (1-7b) of the first row represent positions on the premaxilla Each row corresponds to an individual stickleback larva, with colored boxes showing the tooth sequence for each corresponding fish. Colors/shades indicate stage of tooth at that position: absent (light gray); developing tooth (green); ankylosed tooth (dark). Dpf = days post fertilization (n=76).

**Figure S3.**

| Time Course | Fish ID | Reference | dpf | Freshwater Premaxilla 1 | 2 | 3 | 4 | 5 | 6 | 7a | 7b | 7c |
|---|---|---|---|---|---|---|---|---|---|---|---|---|
| TC4 | 10-2 | 1 | 10 | | | | | | | | | |
| TC4 | 10-4 | 2 | 10 | | | | | | | | | |
| TC4 | 11-2 | 3 | 11 | | | | | | | | | |
| TC4 | 11-3 | 4 | 11 | | | | | | | | | |
| TC4 | 11-4 | 5 | 11 | | | | | | | | | |
| TC6 | 12-1 | 6 | 11 | | | | | | | | | |
| TC6 | 12-2 | 7 | 11 | | | | | | | | | |
| TC2 | 14-2 | 8 | 12 | | | | | | | | | |
| TC2 | 14-3 | 9 | 12 | | | | | | | | | |
| TC4 | 12-1 | 10 | 12 | | | | | | | | | |
| TC4 | 12-2 | 11 | 12 | | | | | | | | | |
| TC4 | 12-3 | 12 | 12 | | | | | | | | | |
| TC4 | 12-4 | 13 | 12 | | | | | | | | | |
| TC5 | 14-1 | 14 | 12 | | | | | | | | | |
| TC5 | 14-2 | 15 | 12 | | | | | | | | | |
| TC5 | 14-3 | 16 | 12 | | | | | | | | | |
| TC5 | 14-4 | 17 | 12 | | | | | | | | | |
| TC6 | 13-1 | 18 | 12 | | | | | | | | | |
| TC6 | 13-2 | 19 | 12 | | | | | | | | | |
| TC6 | 13-3 | 20 | 12 | | | | | | | | | |
| TC6 | 13-4 | 21 | 12 | | | | | | | | | |
| TC6 | 13-5 | 22 | 12 | | | | | | | | | |
| TC6 | 13-6 | 23 | 12 | | | | | | | | | |
| TC3 | 13-1 | 24 | 13 | | | | | | | | | |
| TC3 | 13-2 | 25 | 13 | | | | | | | | | |
| TC3 | 13-3 | 26 | 13 | | | | | | | | | |
| TC3 | 13-4 | 27 | 13 | | | | | | | | | |
| TC4 | 13-1 | 28 | 13 | | | | | | | | | |
| TC4 | 13-2 | 29 | 13 | | | | | | | | | |
| TC4 | 13-3 | 30 | 13 | | | | | | | | | |
| TC4 | 13-4 | 31 | 13 | | | | | | | | | |
| TC2 | 16-3 | 32 | 14 | | | | | | | | | |
| TC2 | 16-4 | 33 | 14 | | | | | | | | | |
| TC4 | 14-1 | 34 | 14 | | | | | | | | | |
| TC4 | 14-3 | 35 | 14 | | | | | | | | | |
| TC4 | 14-4 | 36 | 14 | | | | | | | | | |
| TC5 | 16-1 | 37 | 14 | | | | | | | | | |
| TC5 | 16-2 | 38 | 14 | | | | | | | | | |
| TC5 | 16-3 | 39 | 14 | | | | | | | | | |
| TC5 | 16-4 | 40 | 14 | | | | | | | | | |
| TC3 | 15-1 | 41 | 15 | | | | | | | | | |
| TC3 | 15-2 | 42 | 15 | | | | | | | | | |
| TC3 | 15-3 | 43 | 15 | | | | | | | | | |
| TC3 | 15-4 | 44 | 15 | | | | | | | | | |
| TC3 | 15-5 | 45 | 15 | | | | | | | | | |
| TC4 | 15-1 | 46 | 15 | | | | | | | | | |
| TC4 | 15-2 | 47 | 15 | | | | | | | | | |
| TC4 | 15-3 | 48 | 15 | | | | | | | | | |
| TC4 | 15-4 | 49 | 15 | | | | | | | | | |
| TC2 | 18-3 | 50 | 16 | | | | | | | | | |
| TC4 | 16-1 | 51 | 16 | | | | | | | | | |
| TC4 | 16-2 | 52 | 16 | | | | | | | | | |
| TC4 | 16-3 | 53 | 16 | | | | | | | | | |
| TC5 | 18-1 | 54 | 16 | | | | | | | | | |
| TC5 | 18-2 | 55 | 16 | | | | | | | | | |
| TC5 | 18-3 | 56 | 16 | | | | | | | | | |
| TC3 | 17-1 | 57 | 17 | | | | | | | | | |
| TC3 | 17-3 | 58 | 17 | | | | | | | | | |
| TC4 | 18-1 | 59 | 18 | | | | | | | | | |
| TC4 | 18-2 | 60 | 18 | | | | | | | | | |
| TC4 | 18-3 | 61 | 18 | | | | | | | | | |
| TC2 | 22-1 | 62 | 20 | | | | | | | | | |
| TC5 | 22-2 | 63 | 20 | | | | | | | | | |
| TC4 | 25-2 | 64 | 25 | | | | | | | | | |
| TC4 | 25-3 | 65 | 25 | | | | | | | | | |
| TC5 | 28-1 | 66 | 26 | | | | | | | | | |
| TC5 | 28-3 | 67 | 26 | | | | | | | | | |
| TC5 | 28-4 | 68 | 26 | | | | | | | | | |
| TC5 | 30-1 | 69 | 28 | | | | | | | | | |
| TC5 | 30-2 | 70 | 28 | | | | | | | | | |
| TC5 | 30-3 | 71 | 28 | | | | | | | | | |
| TC5 | 30-4 | 72 | 28 | | | | | | | | | |
| TC5 | 30-5 | 73 | 28 | | | | | | | | | |

Developing Germ

Ankylosed Tooth

Absent

**Figure S3. Scoring of the primary premaxillary tooth sequence in freshwater larvae.** Numbers (1-7b) of the first row represent positions on the premaxilla. Each row corresponds to an individual stickleback larva, with colored boxes showing the tooth sequence for each corresponding fish. Colors/shades indicate stage of tooth at that position: absent (light gray); developing tooth (green); ankylosed tooth (dark). Dpf = days post fertilization (n=73).

#### Figure S4.

| Marine Dentary | | | | Pioneer | Medial | Lateral | | | | | | Developing Germ |
|---|---|---|---|---|---|---|---|---|---|---|---|---|
| Time Course | Fish ID | Reference | dpf | 1 | 2 | 3 | 4 | 5 | 6 | 7 | 8 | Ankylosed Tooth |
| TC5 | 9-1 | 1 | 9 | | | | | | | | | Absent |
| TC5 | 9-2 | 2 | 9 | | | | | | | | | |
| TC5 | 9-3 | 3 | 9 | | | | | | | | | |
| TC5 | 9-4 | 4 | 9 | | | | | | | | | |
| TC5 | 10-1 | 5 | 10 | | | | | | | | | |
| TC5 | 10-2 | 6 | 10 | | | | | | | | | |
| TC5 | 10-3 | 7 | 10 | | | | | | | | | |
| TC5 | 10-4 | 8 | 10 | | | | | | | | | |
| TC3 | 11-1 | 9 | 11 | | | | | | | | | |
| TC3 | 11-3 | 10 | 11 | | | | | | | | | |
| TC4 | 10-2 | 11 | 11 | | | | | | | | | |
| TC4 | 10-3 | 12 | 11 | | | | | | | | | |
| TC3 | 12-1 | 13 | 12 | | | | | | | | | |
| TC3 | 12-2 | 14 | 12 | | | | | | | | | |
| TC4 | 11-1 | 15 | 12 | | | | | | | | | |
| TC6 | 12-1 | 16 | 12 | | | | | | | | | |
| TC6 | 12-2 | 17 | 12 | | | | | | | | | |
| TC6 | 12-3 | 18 | 12 | | | | | | | | | |
| TC6 | 12-4 | 19 | 12 | | | | | | | | | |
| TC6 | 12-5 | 20 | 12 | | | | | | | | | |
| TC6 | 12-6 | 21 | 12 | | | | | | | | | |
| TC6 | 12-8 | 22 | 12 | | | | | | | | | |
| TC6 | 12-10 | 23 | 12 | | | | | | | | | |
| TC3 | 13-1 | 24 | 13 | | | | | | | | | |
| TC4 | 12-2 | 25 | 13 | | | | | | | | | |
| TC4 | 12-3 | 26 | 13 | | | | | | | | | |
| TC4 | 12-4 | 27 | 13 | | | | | | | | | |
| TC5 | 13-1 | 28 | 13 | | | | | | | | | |
| TC5 | 13-2 | 29 | 13 | | | | | | | | | |
| TC5 | 13-3 | 30 | 13 | | | | | | | | | |
| TC6 | 13-2 | 31 | 13 | | | | | | | | | |
| TC6 | 13-3 | 32 | 13 | | | | | | | | | |
| TC6 | 13-9 | 33 | 13 | | | | | | | | | |
| TC6 | 13-10 | 34 | 13 | | | | | | | | | |
| TC2 | 16-1 | 35 | 14 | | | | | | | | | |
| TC2 | 16-2 | 36 | 14 | | | | | | | | | |
| TC2 | 16-3 | 37 | 14 | | | | | | | | | |
| TC4 | 13-2 | 38 | 14 | | | | | | | | | |
| TC3 | 15-3 | 39 | 15 | | | | | | | | | |
| TC4 | 14-1 | 40 | 15 | | | | | | | | | |
| TC4 | 14-3 | 41 | 15 | | | | | | | | | |
| TC2 | 18-3 | 42 | 16 | | | | | | | | | |
| TC4 | 15-1 | 43 | 16 | | | | | | | | | |
| TC3 | 17-1 | 44 | 17 | | | | | | | | | |
| TC3 | 17-2 | 45 | 17 | | | | | | | | | |
| TC5 | 18-1 | 46 | 18 | | | | | | | | | |
| TC5 | 18-2 | 47 | 18 | | | | | | | | | |
| TC5 | 18-3 | 48 | 18 | | | | | | | | | |
| TC5 | 18-4 | 49 | 18 | | | | | | | | | |
| TC3 | 19-1 | 50 | 19 | | | | | | | | | |
| TC5 | 19-1 | 51 | 19 | | | | | | | | | |
| TC5 | 19-2 | 52 | 19 | | | | | | | | | |
| TC5 | 19-3 | 53 | 19 | | | | | | | | | |
| TC5 | 19-4 | 54 | 19 | | | | | | | | | |
| TC2 | 22-2 | 55 | 20 | | | | | | | | | |
| TC5 | 20-1 | 56 | 20 | | | | | | | | | |
| TC5 | 20-2 | 57 | 20 | | | | | | | | | |
| TC5 | 20-3 | 58 | 20 | | | | | | | | | |
| TC5 | 20-4 | 59 | 20 | | | | | | | | | |
| TC3 | 21-1 | 60 | 21 | | | | | | | | | |
| TC5 | 25-1 | 61 | 25 | | | | | | | | | |
| TC5 | 25-2 | 62 | 25 | | | | | | | | | |
| TC5 | 25-3 | 63 | 25 | | | | | | | | | |
| TC5 | 25-4 | 64 | 25 | | | | | | | | | |
| TC5 | 26-1 | 65 | 26 | | | | | | | | | |
| TC5 | 26-2 | 66 | 26 | | | | | | | | | |
| TC5 | 26-3 | 67 | 26 | | | | | | | | | |
| TC5 | 26-4 | 68 | 26 | | | | | | | | | |
| TC5 | 27-2 | 69 | 27 | | | | | | | | | |
| TC5 | 27-4 | 70 | 27 | | | | | | | | | |
| TC5 | 28-3 | 71 | 28 | | | | | | | | | |
| TC5 | 28-4 | 72 | 28 | | | | | | | | | |

**Figure S4. Scoring of the primary dentary tooth sequence in marine larvae.** Numbers (1-7b) of the first row represent positions on the dentary. Each row corresponds to an individual stickleback larva, with colored boxes showing the tooth sequence for each corresponding fish. Colors/shades indicate stage of tooth at that position: absent (light gray); developing tooth (green); ankylosed tooth (dark). Dpf = days post fertilization (n=72).

**Figure S5.**

| Time Course | Fish ID | Freshwater Dentary Reference | dpf | Pioneer 1 | Medial 2 | Lateral 3 | 4 | 5 | 6 | 7 | 8 |
|---|---|---|---|---|---|---|---|---|---|---|---|
| TC4 | 10-2 | 1 | 10 | | | | | | | | |
| TC4 | 10-4 | 2 | 10 | | | | | | | | |
| TC4 | 11-2 | 3 | 11 | | | | | | | | |
| TC4 | 11-3 | 4 | 11 | | | | | | | | |
| TC4 | 11-4 | 5 | 11 | | | | | | | | |
| TC6 | 12-1 | 6 | 11 | | | | | | | | |
| TC6 | 12-2 | 7 | 11 | | | | | | | | |
| TC2 | 14-2 | 8 | 12 | | | | | | | | |
| TC2 | 14-3 | 9 | 12 | | | | | | | | |
| TC4 | 12-1 | 10 | 12 | | | | | | | | |
| TC4 | 12-2 | 11 | 12 | | | | | | | | |
| TC4 | 12-3 | 12 | 12 | | | | | | | | |
| TC4 | 12-4 | 13 | 12 | | | | | | | | |
| TC5 | 14-1 | 14 | 12 | | | | | | | | |
| TC5 | 14-2 | 15 | 12 | | | | | | | | |
| TC5 | 14-3 | 16 | 12 | | | | | | | | |
| TC5 | 14-4 | 17 | 12 | | | | | | | | |
| TC6 | 13-1 | 18 | 12 | | | | | | | | |
| TC6 | 13-2 | 19 | 12 | | | | | | | | |
| TC6 | 13-3 | 20 | 12 | | | | | | | | |
| TC6 | 13-4 | 21 | 12 | | | | | | | | |
| TC6 | 13-5 | 22 | 12 | | | | | | | | |
| TC6 | 13-6 | 23 | 12 | | | | | | | | |
| TC6 | 13-7 | 24 | 12 | | | | | | | | |
| TC3 | 13-1 | 25 | 13 | | | | | | | | |
| TC3 | 13-2 | 26 | 13 | | | | | | | | |
| TC3 | 13-3 | 27 | 13 | | | | | | | | |
| TC3 | 13-4 | 28 | 13 | | | | | | | | |
| TC4 | 13-1 | 29 | 13 | | | | | | | | |
| TC4 | 13-2 | 30 | 13 | | | | | | | | |
| TC4 | 13-3 | 31 | 13 | | | | | | | | |
| TC4 | 13-4 | 32 | 13 | | | | | | | | |
| TC2 | 16-3 | 33 | 14 | | | | | | | | |
| TC2 | 16-4 | 34 | 14 | | | | | | | | |
| TC4 | 14-1 | 35 | 14 | | | | | | | | |
| TC4 | 14-3 | 36 | 14 | | | | | | | | |
| TC4 | 14-4 | 37 | 14 | | | | | | | | |
| TC5 | 16-1 | 38 | 14 | | | | | | | | |
| TC5 | 16-2 | 39 | 14 | | | | | | | | |
| TC5 | 16-3 | 40 | 14 | | | | | | | | |
| TC5 | 16-4 | 41 | 14 | | | | | | | | |
| TC3 | 15-1 | 42 | 15 | | | | | | | | |
| TC3 | 15-2 | 43 | 15 | | | | | | | | |
| TC3 | 15-3 | 44 | 15 | | | | | | | | |
| TC3 | 15-4 | 45 | 15 | | | | | | | | |
| TC3 | 15-5 | 46 | 15 | | | | | | | | |
| TC4 | 15-1 | 47 | 15 | | | | | | | | |
| TC4 | 15-2 | 48 | 15 | | | | | | | | |
| TC4 | 15-3 | 49 | 15 | | | | | | | | |
| TC4 | 15-4 | 50 | 15 | | | | | | | | |
| TC2 | 18-2 | 51 | 16 | | | | | | | | |
| TC4 | 16-1 | 52 | 16 | | | | | | | | |
| TC4 | 16-2 | 53 | 16 | | | | | | | | |
| TC4 | 16-3 | 54 | 16 | | | | | | | | |
| TC5 | 18-1 | 55 | 16 | | | | | | | | |
| TC5 | 18-2 | 56 | 16 | | | | | | | | |
| TC5 | 18-3 | 57 | 16 | | | | | | | | |
| TC3 | 17-1 | 58 | 17 | | | | | | | | |
| TC3 | 17-3 | 59 | 17 | | | | | | | | |
| TC2 | 20-1 | 60 | 18 | | | | | | | | |
| TC4 | 18-1 | 61 | 18 | | | | | | | | |
| TC4 | 18-2 | 62 | 18 | | | | | | | | |
| TC4 | 18-3 | 63 | 18 | | | | | | | | |
| TC2 | 22-1 | 64 | 20 | | | | | | | | |
| TC5 | 22-1 | 65 | 20 | | | | | | | | |
| TC5 | 22-2 | 66 | 20 | | | | | | | | |
| TC5 | 22-3 | 67 | 20 | | | | | | | | |
| TC5 | 22-4 | 68 | 20 | | | | | | | | |
| TC4 | 25-1 | 69 | 25 | | | | | | | | |
| TC4 | 25-2 | 70 | 25 | | | | | | | | |
| TC4 | 25-3 | 71 | 25 | | | | | | | | |
| TC5 | 28-1 | 72 | 26 | | | | | | | | |
| TC5 | 28-2 | 73 | 26 | | | | | | | | |
| TC5 | 28-3 | 74 | 26 | | | | | | | | |
| TC5 | 28-4 | 75 | 26 | | | | | | | | |
| TC5 | 30-1 | 76 | 28 | | | | | | | | |
| TC5 | 30-2 | 77 | 28 | | | | | | | | |
| TC5 | 30-3 | 78 | 28 | | | | | | | | |
| TC5 | 30-4 | 79 | 28 | | | | | | | | |
| TC5 | 30-5 | 80 | 28 | | | | | | | | |

Developing Germ
Ankylosed Tooth
**Absent**

**Figure S5. Scoring of the primary dentary tooth sequence in freshwater larvae.** Numbers (1-7b) of the first row represent positions on the dentary. Each row corresponds to an individual stickleback larva, with colored boxes showing the tooth sequence for each corresponding fish. Colors/shades indicate stage of tooth at that position: absent (light gray); developing tooth (green); ankylosed tooth (dark). Dpf = days post fertilization (n=80).

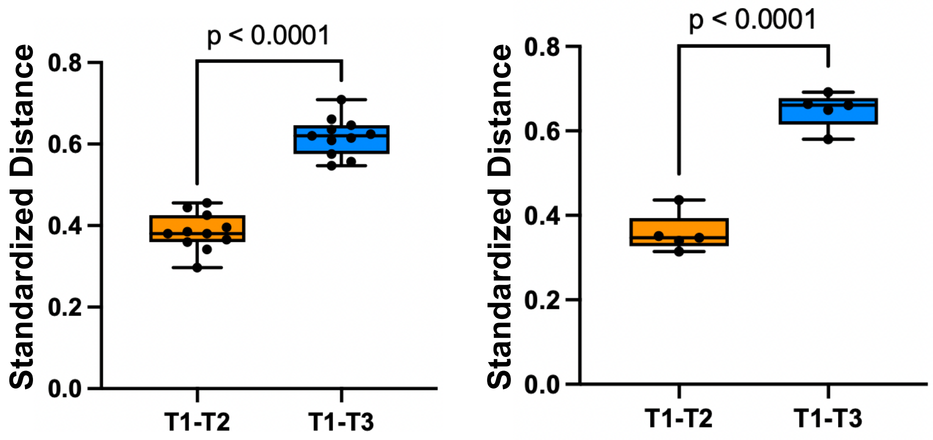


**Figure S6. Significant difference in the spacing between the pioneer tooth and the tooth in the medial and lateral positions on the premaxilla.** Standardized distance between the pioneer tooth and the adjacent tooth in the medial and lateral position in (A) marine and (B) freshwater populations. T1=Pioneer tooth, T2=medial position, T3=lateral position (Total n=16). Standardized distance: 1= 100µm.

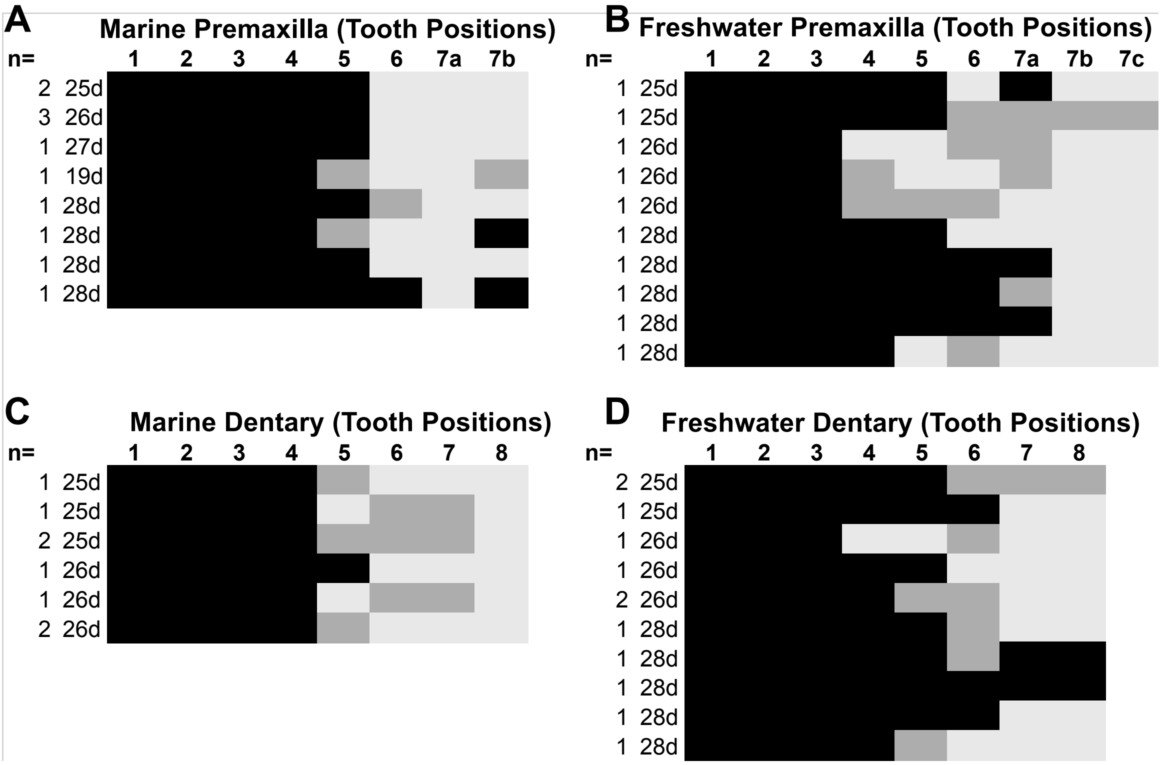


**Figure S7. Variability in premaxillary and dentary tooth sequence increases after the four-tooth stage.** Tooth sequences of marine and freshwater fish between 25-28 dpf (days post-fertilization). (A-B) Tooth positions occupied on the premaxilla of (A) marine and (B) freshwater populations. (B-C) Dentary tooth positions in (C) marine and (D) freshwater fish. Absent (light gray); developing tooth (dark gray); ankylosed tooth (black). n = 23 (marine = 11; freshwater = 12).

Figure S8.

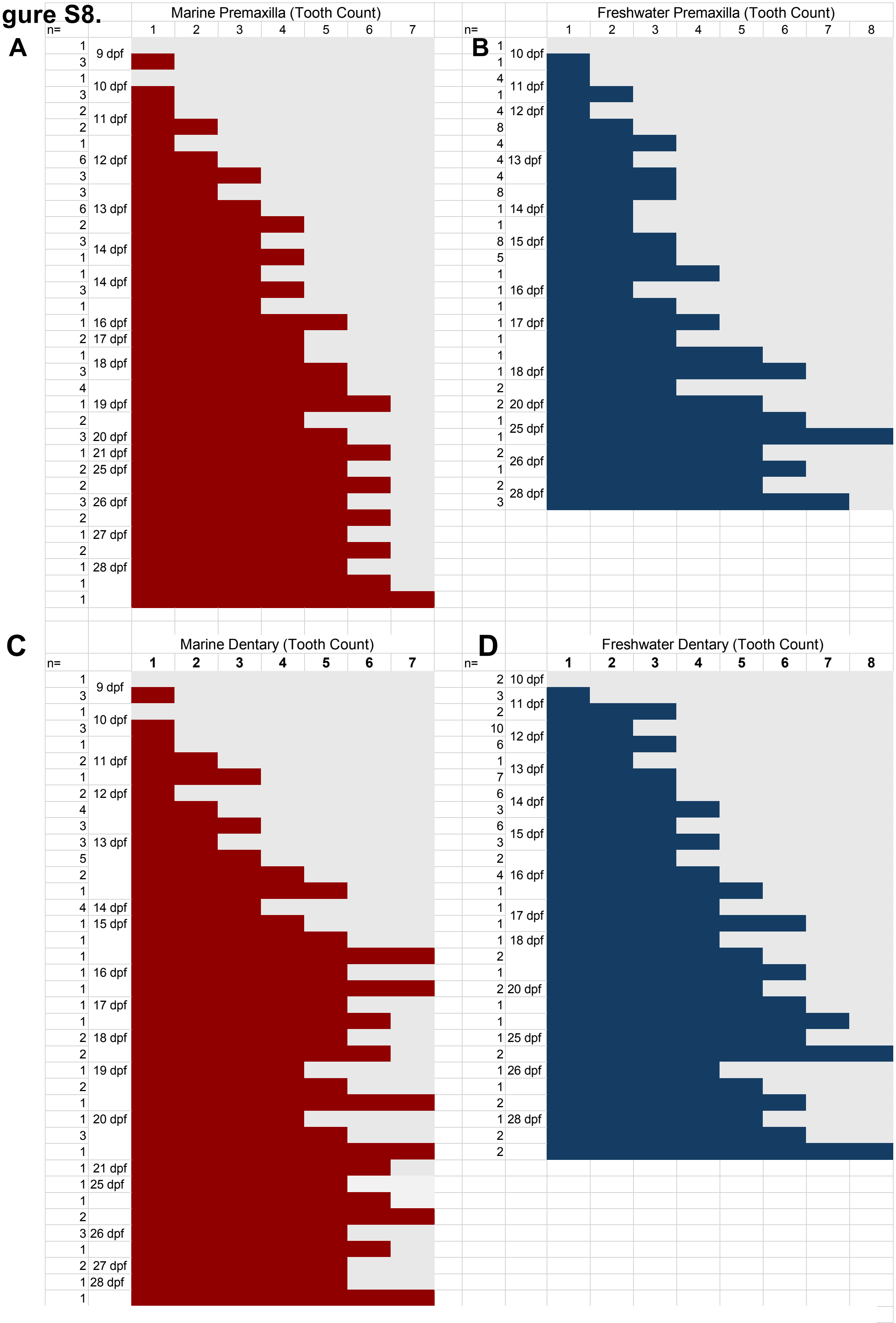
A
Marine Premaxilla (Tooth Count)
n=
1 2 3 4 5 6 7
1 3 1 3 2 2 1 6 3 3 6 2 3 1 1 3 1 1 2 1 3 4 1 2 3 1 2 2 3 2 1 2 1 1 1
9 dpf 10 dpf 11 dpf 12 dpf 13 dpf 14 dpf 14 dpf 16 dpf 17 dpf 18 dpf 19 dpf 20 dpf 21 dpf 25 dpf 26 dpf 27 dpf 28 dpf
B
Freshwater Premaxilla (Tooth Count)
n=
1 2 3 4 5 6 7 8
1 1 4 1 4 8 4 4 4 8 1 1 8 5 1 1 1 1 1 1 1 2 2 1 1 2 1 2 3
10 dpf 11 dpf 12 dpf 13 dpf 14 dpf 15 dpf 16 dpf 17 dpf 18 dpf 20 dpf 25 dpf 26 dpf 28 dpf
C
Marine Dentary (Tooth Count)
n=
1 2 3 4 5 6 7
1 3 1 3 1 2 1 2 4 3 3 5 2 1 4 1 1 1 1 1 1 1 2 2 1 2 1 1 3 1 1 1 1 2 3 1 2 1 1
9 dpf 10 dpf 11 dpf 12 dpf 13 dpf 14 dpf 15 dpf 16 dpf 17 dpf 18 dpf 19 dpf 20 dpf 21 dpf 25 dpf 26 dpf 27 dpf 28 dpf
D
Freshwater Dentary (Tooth Count)
n=
1 2 3 4 5 6 7 8
2 3 2 10 6 1 7 6 3 6 3 2 4 1 1 1 1 2 1 2 1 1 1 2 1 1 2 1 2 2
10 dpf 11 dpf 12 dpf 13 dpf 14 dpf 15 dpf 16 dpf 17 dpf 18 dpf 20 dpf 25 dpf 26 dpf 28 dpf

**Figure S8. Oral tooth counts in marine and freshwater fish early in development.** (A-B) Total premaxillary tooth counts in (A) marine and (B) freshwater sticklebacks from 9-28 dpf. (C-D) Total dentary tooth count in (C) marine and (D) freshwater sticklebacks. Numbered columns indicate the tooth count present in individual fish. Horizontal rows indicate the days post fertilization (dpf) at which that tooth count was present. Marine n = 75 (premaxilla); n=69 (dentary); freshwater: n=75 (premaxilla); n=78 (dentary).

| | | | Resorption Crater | | | Multi-rowed Dentition | |
|---|---|---|---|---|---|---|---|
| Fish ID | dpf | SL (mm) | Premaxilla | Dentary | | Premaxilla | Dentary |
| 080525-1 | 40d | 10.96 | | | | | |
| 080525-2 | 40d | 8.34 | | | | | |
| 080525-3 | 40d | 10.57 | | | | | |
| 080625-1 | 41d | 9.37 | | | | | |
| 080625-2 | 41d | 9.5 | | | | | |
| 080625-3 | 41d | 11.28 | | | | | |
| 080625-4 | 41d | 10.01 | | | | | |
| 080825-1 | 43d | 11.24 | | | | | |
| 080825-2 | 43d | 9.87 | | | | | |
| 080825-3 | 43d | 13.47 | | | | | |

**Figure S9. Scoring of morphological indications of tooth shedding and replacement.** Morphological indicators: (1) resorption craters and (2) multi-rowed dentition in freshwater sticklebacks. Shading of rectangular boxes indicate whether the morphological observation was present (dark grey) or absent (light grey) in that jaw for each fish. dpf = days post fertilization. SL = standard length (mm).
